# Error sensitivity and optimization of steady-state kinetic parameters using multidimensional chemical kinetic analysis

**DOI:** 10.1101/474171

**Authors:** Joseph S. Eskew, Christopher G. Connell, Jared C. Cochran

**Affiliations:** Department of Molecular & Cellular Biochemistry, Indiana University, Bloomington, IN 47405, USA.; Department of Mathematics, Indiana University, Bloomington, IN 47405, USA.

## Abstract

Enzyme behavior has been described using the Michaelis-Menten mechanism. The analysis of extended time domains provides a means to extract the Michaelis-Menten constants through direct fitting of raw data. We have developed a scheme for determining Michaelis-Menten rate constants by appropriate fitting of multidimensional experimental data sets to the closed form of the Michaelis-Menten model. We considered how varying parameters in experimental data affect the accuracy of the remaining parameter estimates. We determine how to improve experimental design to achieve a given accuracy, relative to the amount of intrinsic or external error. We analyze this scheme on data sets built around 20 hypothetical and 2 natural enzymes (kinesin and apyrase) to test error sensitivity in different parameter regimes. Overall, we provide evidence that our data fitting regime will tolerate significant experimental error in the raw data and still converge on the four Michaelis-Menten constants.

## Introduction

Enzymes are proteins that catalyze specific biochemical reactions in living cells to generate needed products for metabolic and/or replicative processes. Scientists have been studying enzymes for decades and a wealth of information is available for many different types of enzymes that exist in all domains of life (Eukarya, Bacteria, and Archaea). The Michaelis-Menten (MM)^1^ mechanism has been the foundation for studying the steady-state kinetic behavior of enzymatic reactions for over 100 years [1]. In the basic MM scheme (Equation 1), a substrate (S) binds reversibly with a free enzyme (E) to form a non-covalent enzyme-substrate complex, which reacts irreversibly to yield free product (P) and the regenerated free enzyme. One can easily demonstrate the hyperbolic relationship between the initial reaction velocity and substrate concentration through mathematical derivation under a given set of basic assumptions [2]. However, these assumptions are not always valid for enzymatic systems and/or experimental design regimens. This is especially true for extended time domains that allow for both substrate exhaustion and product inhibition of enzyme activity through binding competition. Therefore, more complex mechanisms must be employed to consider the biochemical steps for chemistry (ES ⇌ EP) and product release and re-binding to free enzyme (EP ⇌ E + P).

MM steady-state experiments focus on capturing data for product formation during the initial velocity time regime with variant substrate concentrations. These raw data are subsequently fit to a linear function and the slopes that define the initial reaction velocities are plotted against the substrate concentration to be fit to the hyperbolic MM equation. This process leads to extraction of the *k*_*cat*_ and *K*_*S*_ (also called the *K*_*m*_) parameter for the enzyme-catalyzed reaction. Due to restrictions that are directly related to the experimental methodologies (e.g. linear range for product detection) or to the nature of the enzyme under investigation (e.g. an enzyme that binds its substrate very tightly but has a very slow maximal rate of substrate turnover), these experiments often do not accurately define the intrinsic rate and equilibrium constants. Also, when these experiments do work as intended, the enzymologist would only yield two of the multiple parameters that govern the overall enzymatic cycle.

A real challenge in obtaining reliable steady state values centers on designing experiments to minimize error and maximize information about the enzyme mechanism. Since results from steady-state experiments are typically the first insight into the mechanistic character of an unknown enzyme, having a more thorough analysis gives the enzymologist a better understanding for designing more complicated kinetic and/or thermodynamic experiments. Obtaining microscopic rate constants are typically validated by numerous orthogonal chemical and physical studies, such as isotope and viscosity effects [3–9]. More elaborate single turnover and transient state kinetic experiments can be employed to define intrinsic rate constants in the minimal enzyme mechanism [10–15]. Together, data from multiple kinetic approaches can be globally fit based on numerical integration of the rate equations using a host of kinetic simulation software packages including KINSIM/FITSIM [16, 17], DYNAFIT [18], COPASI [6, 19], KinTek Global Explorer [20, 21], and Berkeley Madonna™. Current computation speed has made these algorithms very effective, especially in one and two dimensions, for recovering best-fits for rate constants from a given sampling of data without the need to explicitly solve the underlying model. However, as the number of simultaneously unknown rate constants increases, corresponding to the dimension of numerical integration, e.g. via the Levenberg–Marquardt algorithm, the relative speed gain from using explicit model solutions increases. Also by their very nature, such integration fitting algorithms are unable to easily exploit the specific features of analytic solutions to achieve the goal of minimizing the data collection needed to obtain reasonable precision of the rate constants. We propose to improve optimized collection through selection of data points based on direct, yet fully automated, analysis of features in analytical solutions of the specific model. This allows us to achieve surprising accuracy in fitting even three or more rate constants simultaneously from a sparing number of experimental data points.

Since the seminal work of MM [22], reaction kinetics have been monitored beyond the initial linear phase into a phase where substrate deletion and product accumulation results in slowing the instantaneous velocity of the reaction, approaching a velocity of zero at infinite time (or at equilibrium). The analysis of these long time domains through fitting of progress curves has provided an alternative method for ascertaining the parameters that govern the basic MM steady-state mechanism [23–27]. Nevertheless, the sensitivity of the fitting regimes has not been well defined [28], especially along the axes for experimental conditions that can be varied for any given enzymatic system. We attempt to address these issues with a fitting scheme outlined in this manuscript. We analyze for accuracy in recovering known rate constants from data in the presence of various error regimes. Parameter redundancy is elucidated explicitly, allowing us to both avoid redundant sampling on the one hand, and to test its effects on poorly chosen data as well. Several statistical comparisons of the efficiency of different experimental designs are performed. These especially focus on contrasting the effectiveness of finding unknown parameters starting from different known parameters and how sample size and the dimension of the data collected affects accuracy.

## Materials and Methods

### Experimental conditions and proteins used

The DNA construct coding for the motor domain of human kinesin-5 (Eg5; residues 1–368) was cloned into a modified pET16b (Novagen) vector for bacterial expression to achieve an N-terminally 6x His tag and TEV cleavage sequence. (N-terminus: MGSSHHHHHH-SSGENLYFQGSH-^1^MASQ). Expression and purification of nucleotide-free Eg5 was performed as previously described [29–32]. Kinetic experiments for Eg5 were performed in assay buffer 1 (20 mM HEPES pH 7.2 with KOH, 5 mM magnesium acetate, 0.1 mM EGTA, 0.1 mM EDTA, 50 mM potassium acetate, 150 mM sucrose, 1 mM DTT) at 298 K as described previously [30, 31, 33]. Microtubules (MTs) were prepared using an aliquot of purified bovine brain tubulin that was thawed and cycled, and the MTs were stabilized with 20 μM paclitaxel (Sigma-Aldrich). Kinetic experiments for *Solanum tuberosum* apyrase (Apy; Grade VII; Sigma-Aldrich) were performed in assay buffer 2 (10 mM HEPES pH 7.2 with KOH, 15 mM MgCl_2_, 1 mM EGTA, 100 mM NaCl, 150 mM sucrose, 0.02% v/v TWEEN 20) at 298 K with the concentrations reported as final after mixing. Protein concentrations were determined using the Bradford assay (Sigma-Aldrich) with bovine serum albumin as standard.

### Kinetic assays

For microtubule-stimulated ATPase measurements, Eg5 at 0.2 μM was complexed with paclitaxel-stabilized microtubules (MTs) at 2 μM and reacted with varying concentrations of ATP substrate (95–310 μM) and added ADP product (0 or 1500 μM). Samples of the reaction (10 μL each) were quenched in 1 N HCl (2.5 μL of 5 N stock) followed by addition of malachite green reagent (150 μL; room temperature; [34]). After thorough mixing by vortexing, 150 μL of this mixture was transferred to a 96-well plate for endpoint absorbance (λ=650 nm) using a BioTek Epoch microplate spectrophotometer. A standard curve from 10–400 μM Na_2_HPO_4_/NaH_2_PO_4_ (pH 7.2) was used to convert A_650_ to inorganic phosphate (P_i_) concentration. For each progress curve, P_i_ produced by Eg5 ATPase (total P_i_ minus background P_i_) was plotted against time.

For ADPase measurements, Apy at 0.13 μM was reacted with varying ADP substrate (40–260 μM) and added AMP product (0 or 10 mM). P_i_ product was monitored using the malachite green assay as described above. For each progress curve, P_i_ produced by Apy ADPase was plotted against time. Stock ATP and ADP concentrations were experimentally determined using absorbance (λ=260 nm) with molar extinction coefficient (15,400 L mol^−1^ cm^−1^).

### Mechanisms

Expansions of the Michaelis-Menten (MM) mechanism establish irreversible product formation/release (MM model; Equation 1) as well as competitive product inhibition (PI) of enzyme activity with irreversible chemistry step (PI model; Equation 2) and the fully reversible (FR) three-step mechanism (FR model; Equation 3).

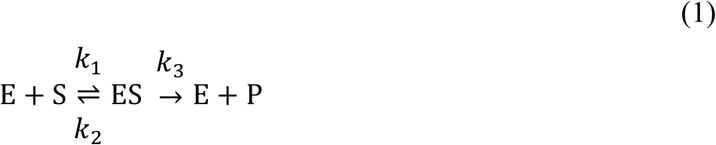

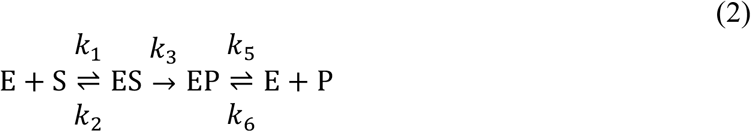

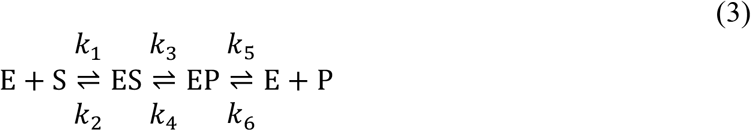

### Closed form of the model

Using *Mathematica* 10 (Wolfram), it was verified by differentiation in time that the differential equation solution takes the closed form of Scheme 1, where *ω* represents the Lambert *ω*-function and *t* is time [35–44].

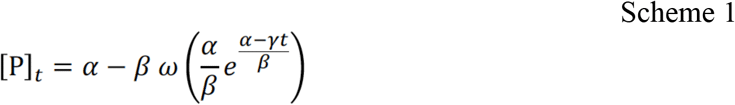

|  | Fully Reversible (FR) | Product Inhibition (PI) | Michaelis-Menten (MM) |
| --- | --- | --- | --- |
| $\alpha$ | $\frac{[S_o]k_{cat}^F K_P - [P_o]k_{cat}^R K_S}{k_{cat}^F K_P + k_{cat}^R K_S}$ | $[S_o]$ | $[S_o]$ |
| $\beta$ | $\frac{K_P K_S \{ [P_o](k_{cat}^R + k_{cat}^F) + [S_o](k_{cat}^R + k_{cat}^F) + k_{cat}^F K_P + k_{cat}^R K_S \}}{(K_P - K_S)(k_{cat}^F K_P + k_{cat}^R K_S)}$ | $\frac{([P_o] + [S_o] + K_P)K_S}{K_P - K_S}$ | $K_S$ |
| $\gamma$ | $[E_o] \frac{k_{cat}^F K_P + k_{cat}^R K_S}{K_P - K_S}$ | $[E_o] \frac{k_{cat}^F K_P}{K_P - K_S}$ | $[E_o]k_{cat}^F$ |

### Asymmetric confidence estimates

Since our simulation analyses tested different magnitudes of total degrees of freedom, we utilized mean squared error (MSE) from nonlinear least squared fitting to construct 3D contour plots for asymmetric confidence estimates based on a similar method of FitSpace Explorer [21]. Briefly, pairs of parameters were fixed and varied around their best fit value, and remaining constants were floated in the fitting scheme. The best fit MSE was divided by the MSE from each fixed pair of constants (MSE_*min*_/MSE_*x,y*_) within a 14×14 matrix centered on the best fit values. Contour plots were constructed in *Mathematica* using these arrays of MSE ratios to linearly interpolate values to provide smooth contours at 0.1 intervals. Bounds of the asymmetric confidence estimates were defined at 11% and 25% increases in MSE (MSE_*min*_/MSE_*x,y*_) = 0.9 red and 0.8 orange, respectively as previously reported [21].

### Error simulation

To generate variably ‘noisy’ data sets that realistically represent a range of experimental observations, normally distributed random additive error (errA) was added to the quantity of product formed during the simulations using Scheme 1 and the following equation:

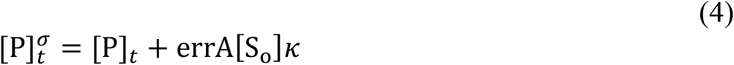

where *κ* represents the fraction of [S_o_] converted to product at equilibrium defined as:

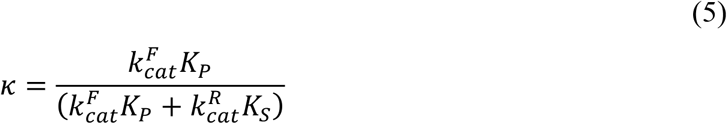

### Time parameter threshold computation

We used the closed form of the model to determine the near-optimal time parameter interval to maximize the efficiency of the fitting algorithms given a fixed sampling frequency. The most valuable portion of the curve is where the absolute value of the second derivative is bounded away from zero, since the remaining portions are nearly linear where the data points are quite redundant for the nonlinear parameter estimation. Toward this end, we compute the derivative of the product formula (Scheme 1). The derivative *ω*′(*t*) a of the Lambert *ω*-function is:

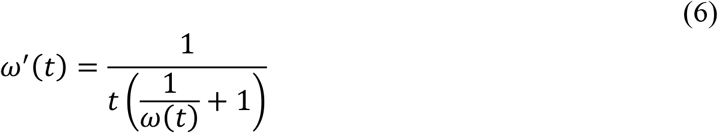

Hence [P]′(*t*)simplifies to:

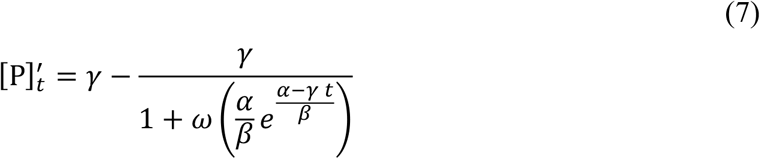

This quantity exponentially decays to 0 as *t* ⟶ ∞. Given this derivative, we chose an appropriate scale to determine when it has become sufficiently small. Since the product [P]_*t*_ is almost exactly linear at the beginning of the time regime, we use the scale of the derivative at 0 to set the appropriate scale.

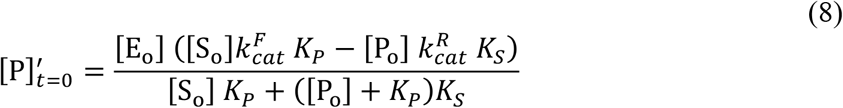

We set the stopping time *t*_*stop*_ to be the time when the derivative reaches 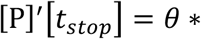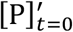 for a given threshold parameter *θ* = *v*_*f*_/*v*_*o*_ where *v*_*f*_ is the instantaneous final velocity and *v*_*o*_ is the initial velocity as *t* ⟶ 0.

### Design for data simulation

Simulated progress curves were generated through calculating the product concentration (Scheme 1) with random error applied (Eqs. 4 and 5) over time under various conditions. SIM1 represents a data set at increasing [S_o_] with no added [P_o_] at *t* = 0. SIM2 represents a data set at constant [S_o_] with increasing [P_o_] that was added at *t* = 0. SIM3 represents a data set at increasing [S_o_] and increasing [P_o_] that was added at *t* = 0. All simulated data sets were designed for equal total degrees of freedom (i.e. equal total data points with 25 points per curve and 9 curves; Table A.1). All statistical evaluations were standard for NonlinearModelFit in *Mathematica*.

### Scoring functions

Judging the quality of the final nonlinear least squares fitting for our SIM1, SIM2, and SIM3 simulations required a scoring function that incorporated both accuracy (i.e. estimated constant from curve fitting divided by the expected constant) and precision (i.e. symmetric 95% confidence interval). In general, more meaningful fitted constants correlate with higher accuracy (~1) and lower confidence intervals. A scoring function was developed to assess each fitted constant:

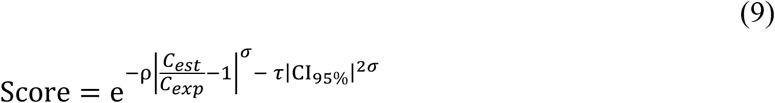

where *C*_*est*_ was the estimated constant from nonlinear least squares fitting, *C*_*exp*_ was the simulated (i.e. expected) constant, and CI_95%_ was the confidence interval (95%) for each measurement based on a simple multiple of the standard error. *ρ*, *σ*, and *τ* are weighting constants chosen to provide a score of 0.5–1 when normalized constants (*C*_*est*_/*C*_*exp*_) are 0.5−1.5 with narrow confidence intervals (< ±50%). For all score calculations presented in this manuscript, *ρ* = 1, *σ* = 0.75, and *τ* = 0.5 (Fig S2*b*).

## Results

### Analysis of experimental kinetic data for Apy and Eg5

Experimental data for kinesin-5 (Eg5) and apyrase (Apy) were fit to either one of three mechanistic models, represented by the algebraic rate equations listed in Scheme 1. The three models, in order of increasing complexity, are designated as “MM”, “PI”, and “FR”, respectively. Model MM represents the simplest Michaelis-Menten kinetic model, whereby the reaction product does not rebind and inhibit the enzyme and the reaction proceeds only in the forward direction (Equation 1). Model PI represents the product inhibition model, whereby the reaction product rebinds and inhibits the enzyme but the overall reaction can only proceed in the forward direction (Equation 2). Model FR represents the fully reversible kinetic model, whereby the reaction product not only rebinds and inhibits the enzyme, but also the overall reaction can proceed in either the forward or reverse direction (Equation 3).

Apy (type VII; EC 3.6.1.5) catalyzes the hydrolysis ADP to AMP and P_i_ [45]. To measure the ADPase kinetics of Apy, we performed a series of reactions at different ADP substrate (40–260 μM) and different AMP product (0 and 10 mM) concentrations and used the malachite green colorimetric assay to quantify P_i_ produced over time. Figs 1*a–b* summarize the results from fitting the Apy ADPase data using basic initial velocity analysis as well as to the three independent models (Table 1). Using both AIC and BIC to assess the goodness of fit relative to the complexity of the model, the apyrase data were best modeled to the FR mechanism (Equation 3) given the lower AIC and BIC values compared to the PI model (ΔAIC = −17 and ΔBIC = −14). However, asymmetric confidence analysis (Table 2; Fig S1*a*) indicated that not all the parameters were well constrained. Specifically, we observed large upper bounds for *K*_*P*_ and 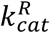 and no lower bound for 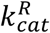. These results suggest that AMP product weakly binds the enzyme yet will proceed in the reverse direction to reform ADP substrate (i.e. the equilibrium constant for the chemistry step was 7 at the best fit values).

**FIGURE 1:**
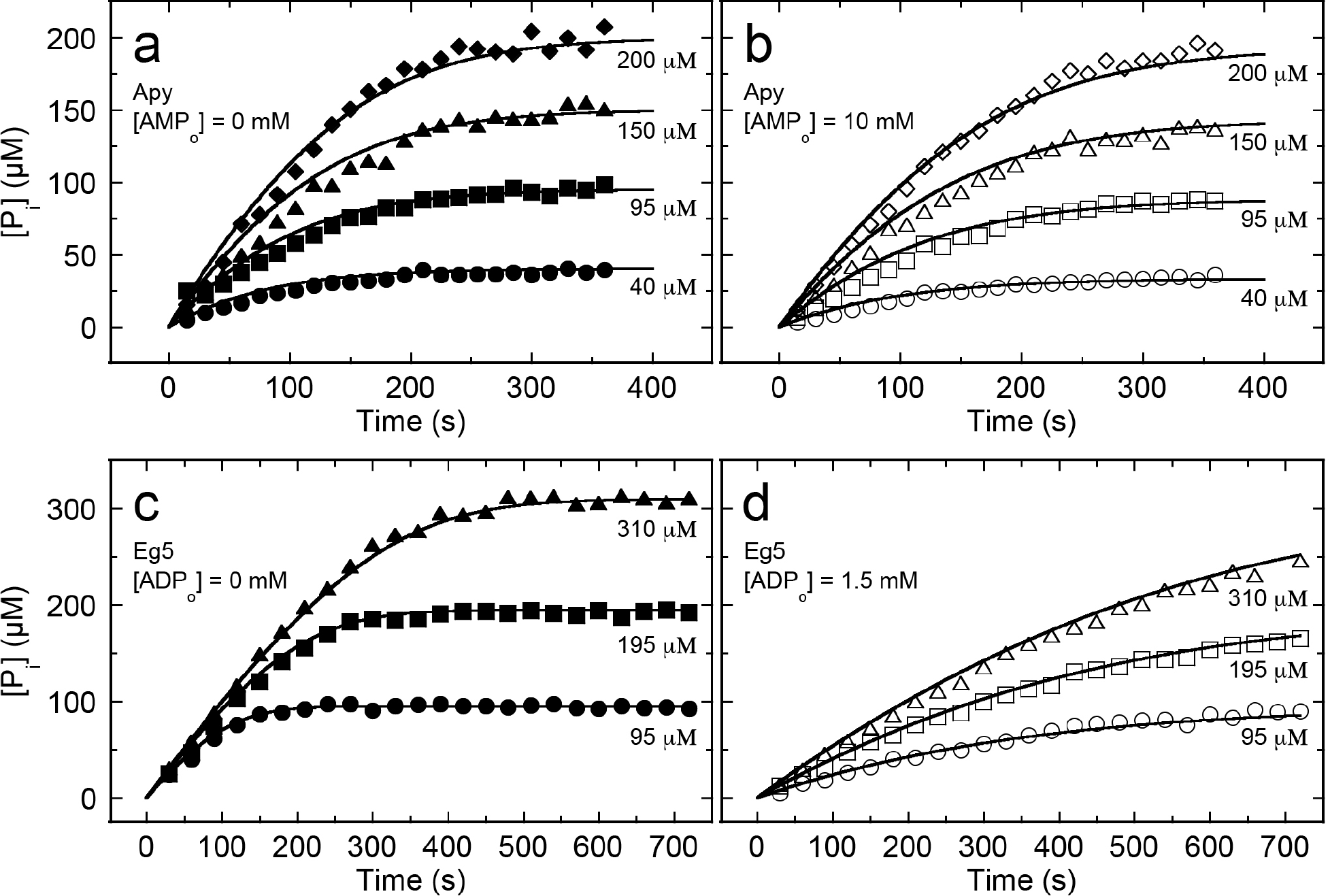
Global fitting of Apy and Eg5 progress curves. (*a*) P_i_ product formation as a function of time for Apy (closed symbols) shown at different initial substrate (ADP) concentrations (as indicated) with no added AMP product at the start of the reactions. (*b*) Time courses of Apy product formation (open symbols) at different substrate concentrations (as indicated) plus added initial AMP product (10 mM) at the start of the reactions, (*c*) P_i_ product formation as a function of time for Eg5 (closed symbols) shown at different initial substrate (ATP) concentrations (as indicated) with no added ADP product at the start of the reactions. (*d*) Time courses of Eg5 product formation (open symbols) at different substrate concentrations (as indicated) plus added initial ADP product (1.5 mM) at the start of the reactions. The smooth lines in each panel represent the best fit global FR model for Apy and PI model for Eg5 using NonlinearModelFit in *Mathematica* with fit constants provided in Table 1.

**Table 1.** Parameter estimates from experimental data for Apy and Eg5.

| Apy<br>Constant | Initial Velocity | MM Model |  | PI Model |  | FR Model |  |
| --- | --- | --- | --- | --- | --- | --- | --- |
| | | Estimate ± SE | CI (95%) | Estimate ± SE | CI (95%) | Estimate ± SE | CI (95%) |
| $K_S$ ($\mu\text{M}$) | $75 \pm 12$ | $153 \pm 22$ | 110–197 | $218 \pm 24$ | 170–266 | $176 \pm 20$ | 137–214 |
| $K_P$ ($\mu\text{M}$) | ND<sup>k</sup> | NA | NA | $25300 \pm 2400$ | 20600–30000 | $38500 \pm 7000$ | 24600–52300 |
| $k_{cat}^F$ ($\text{s}^{-1}$) | $18.7 \pm 0.9$ | $18.3 \pm 1.5$ | 15.3–21.2 | $22.7 \pm 1.6$ | 19.4–25.9 | $19.8 \pm 1.3$ | 17.2–22.4 |
| $k_{cat}^R$ ($\text{s}^{-1}$) | NA | NA | NA | NA | NA | $2.9 \pm 1.1$ | 0.7–5.1 |
| AIC<sup>i</sup> | NA | NA |  | 1620 |  | 1603 |  |
| BIC<sup>j</sup> | NA | NA |  | 1634 |  | 1620 |  |
| Eg5<br>Constant | Initial Velocity<sup>a</sup> | MM Model<sup>b</sup> |  | PI Model<sup>c</sup> |  | FR Model<sup>d</sup> |  |
| | | Estimate ± SE<sup>e</sup> | CI (95%)<sup>f</sup> | Estimate ± SE | CI (95%) | Estimate ± SE | CI (95%) |
| $K_S$ ($\mu\text{M}$) | $7.0 \pm 0.4$<sup>g</sup> | $58.2 \pm 4.6$ | 49–67 | $23.2 \pm 10.3$ | 2.9–43.5 | $23.1 \pm 10.9$ | 2–45 |
| $K_P$ ($\mu\text{M}$) | $117 \pm 86$<sup>g</sup> | NA | NA | $124 \pm 49$ | 28–220 | $124 \pm 49$ | 27–221 |
| $k_{cat}^F$ ($\text{s}^{-1}$) | $5.4 \pm 0.1$<sup>g</sup> | $6.0 \pm 0.1$ | 5.7–6.3 | $5.6 \pm 0.4$ | 4.9–6.3 | $5.6 \pm 0.4$ | 4.8–6.4 |
| $k_{cat}^R$ ($\text{s}^{-1}$) | NA<sup>h</sup> | NA | NA | NA | NA | $1 \times 10^{-13}$ | 0–0.2 |
| AIC | NA | NA |  | 1119 |  | 1121 |  |
| BIC | NA | NA |  | 1131 |  | 1136 |  |

<sup>a</sup> Constants provided by the typical MM analysis of initial velocities. <sup>b</sup> Irreversible MM mechanism defined as in Equation 1. <sup>c</sup> Product inhibition (PI) mechanism defined as in Equation 2. <sup>d</sup> Fully reversible (FR) mechanism as in Equation 3. <sup>e</sup> Nonlinear least-squares fitting estimates with standard error for each fit parameter. <sup>f</sup> CI, confidence interval. <sup>g</sup> Steady-state and equilibrium binding constants obtained under similar experimental conditions [30, 33]. <sup>h</sup> NA, not applicable. <sup>i</sup> AIC, Akaike information criterion. <sup>j</sup> BIC, Bayesian information criterion. <sup>k</sup> ND, not determined.

**Table 2.** Asymmetric confidence limits for Apy and Eg5 experimental data.

| **Apy**<br>Constant | **FR Model**<br>Estimate ± SE | **Confidence**<br>$MSE_{min}/MSE_{x,y} = 0.9$ | **Confidence**<br>$MSE_{min}/MSE_{x,y} = 0.8$ |
| --- | --- | --- | --- |
| $K_S$ ($\mu\text{M}$) | $176 \pm 20$ | 105 – 324 | 83 – 483 |
| $K_P$ ($\mu\text{M}$) | $38,500 \pm 7,000$ | 19,410 – 238,360 | 15,105 – 345,170 |
| $k_{cat}^F$ ($\text{s}^{-1}$) | $19.8 \pm 1.3$ | 16.3 – 32.3 | 14.6 – 43.8 |
| $k_{cat}^R$ ($\text{s}^{-1}$) | $2.9 \pm 1.1$ | 0 – 50 | 0 – 74 |
| **Eg5**<br>Constant | **PI Model**<br>Estimate ± SE | **Confidence**<br>$MSE_{min}/MSE_{x,y} = 0.9$ | **Confidence**<br>$MSE_{min}/MSE_{x,y} = 0.8$ |
| $K_S$ ($\mu\text{M}$) | $23.2 \pm 10.3$ | 0 – 80 | 0 – 134 |
| $K_P$ ($\mu\text{M}$) | $124 \pm 49$ | 0 – 334 | 0 – 457 |
| $k_{cat}^F$ ($\text{s}^{-1}$) | $5.6 \pm 0.4$ | 4.6 – 7.5 | 4.3 – 9.3 |

Eg5 (EC 3.6.4.4) catalyzes the hydrolysis of ATP to ADP and P_i_ products, which can be monitored using the malachite green colorimetric assay [34]. We designed a matrix of microtubule-stimulated Eg5 ATPase reactions with varying ATP substrate (95–310 μM) and added ADP product (0 and 1500 μM) concentrations for our analysis. Figs 1*c–d* summarize the results from fitting the microtubule-stimulated Eg5 ATPase data using the three independent models from above with a direct comparison with previously published results (Table 1) [30, 33]. We found good agreement with the steady-state constants, especially the comparison of the 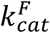 values (5.4 ± 0.1 s^−1^ in previous work [33]; 5.7 ± 0.4 s^−1^ here). The estimates for *K*_*S*_ were consistently 3–8-fold higher than the previously determined MM value, yet the estimate from the PI model was very similar to the *K*_*d,ATP*_ determined from presteady-state acid quench experiments at 20 ± 7 μM [33]. However, increasing the complexity of the model from PI to FR led to a decrease in the goodness of fit with higher AIC and BIC values (ΔAIC = 2 and ΔBIC = 5). These results are consistent with the known chemistry of Eg5, which shows no evidence of ATP synthesis in the presence of microtubules [33]. Asymmetric confidence analysis (Table 2; Fig S1*b*) showed no lower bounds for *K*_*S*_ and *K*_*P*_. Thus, our analysis provides progress curve fitting to extract the binding constants for substrate and product as well as the forward catalytic rate constant with minimal data sets (i.e. 6–8 time courses).

### Simulated enzyme mechanisms

To demonstrate the ability of our multidimensional chemical kinetics analysis of SIM1, SIM2, and SIM3 simulations to extract the four Michaelis-Menten (MM) constants from ‘noisy’ data, we established 2 enzymes based on the best fit constants of Apy and Eg5 as well as 20 hypothetical enzymes that show a wide range of kinetic mechanisms (E1-E20, Table 3, Fig S2*a*). These enzyme mechanisms display 1) widely varying equilibrium constants that span biologically relevant enzymes 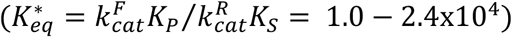, 2) a wide variety of substrate and product binding constants (*K*_*S*_ and *K*_*P*_, respectively) including various combinations seen in natural enzymes, 3) a combinatorial spectrum of forward and reverse catalytic constants 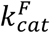 and 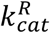, respectively) as well as 4) a range of catalytic efficiencies 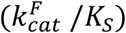 from extremely inefficient enzymes (Apy = 0.13 μM^−1^s^−1^) to a ‘catalytically perfect’ enzyme (E1 = 1003 μM^−1^s^−1^). To generate ‘noisy’ data sets that realistically represent a range of experimental observations, normally distributed random additive error was added to the quantity of product formed during the simulations using Equations 4 and 5 (Fig S3). Given that the four constants 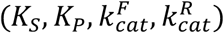 were defined in these simulations, we could assess our nonlinear least squares fitting for both accuracy and precision using a scoring algorithm (Equation 9), which is a function of both the normalized constant (defined as the fitted constant divided by the actual constant) and symmetric 95% confidence interval (Fig S4).

**Table 3.** Simulated enzyme constants.

|  | **Kin5** | **Apy** | **E1** | **E2** | **E3** | **E4** |
| --- | --- | --- | --- | --- | --- | --- |
| $K_S(\mu\text{M})$ | 30 | 150 | 0.11 | 0.11 | 1.1 | 10.1 |
| $K_P(\mu\text{M})$ | 120 | 25000 | 0.12 | 0.12 | 1.2 | 10.2 |
| $k_{cat}^F(\text{s}^{-1})$ | 6 | 20 | 110.3 | 5.3 | 5.3 | 10.3 |
| $k_{cat}^R(\text{s}^{-1})$ | 0.001 | 3 | 2.4 | 0.4 | 0.4 | 0.4 |
| $K_{eq}^*$ | 24000 | 1111 | 50.1 | 14.5 | 14.5 | 26 |
| $k_{cat}^F/K_S(\mu\text{M}^{-1}\text{s}^{-1})$ | 0.2 | 0.13 | 1003 | 48.2 | 4.8 | 1.0 |
|  | **E5** | **E6** | **E7** | **E8** | **E9** | **E10** |
| $K_S(\mu\text{M})$ | 0.11 | 1.1 | 10.1 | 100.1 | 1.1 | 10.1 |
| $K_P(\mu\text{M})$ | 0.12 | 1.2 | 10.2 | 100.2 | 0.62 | 1.2 |
| $k_{cat}^F(\text{s}^{-1})$ | 20.3 | 2.3 | 15.3 | 100.3 | 10.3 | 5.3 |
| $k_{cat}^R(\text{s}^{-1})$ | 20.4 | 2.4 | 15.4 | 100.4 | 4.4 | 0.4 |
| $K_{eq}^*$ | 1.1 | 1.0 | 1.0 | 1.0 | 1.3 | 1.6 |
| $k_{cat}^F/K_S(\mu\text{M}^{-1}\text{s}^{-1})$ | 185 | 2.1 | 1.5 | 1.0 | 9.4 | 0.5 |
|  | **E11** | **E12** | **E13** | **E14** | **E15** | **E16** |
| $K_S(\mu\text{M})$ | 0.1 | 0.1 | 1.1 | 10.1 | 0.1 | 1.1 |
| $K_P(\mu\text{M})$ | 1.2 | 1.2 | 10.2 | 100.2 | 1.2 | 10.2 |
| $k_{cat}^F(\text{s}^{-1})$ | 10.3 | 5.3 | 5.3 | 10.3 | 2.3 | 2.3 |
| $k_{cat}^R(\text{s}^{-1})$ | 4.4 | 0.4 | 0.4 | 3.4 | 2.4 | 2.4 |
| $K_{eq}^*$ | 28.1 | 159 | 123 | 30 | 11.5 | 8.9 |
| $k_{cat}^F/K_S(\mu\text{M}^{-1}\text{s}^{-1})$ | 103 | 53 | 4.8 | 1.0 | 23 | 2.1 |
|  | **E17** | **E18** | **E19** | **E20** |  |  |
| $K_S(\mu\text{M})$ | 1.1 | 10.1 | 0.1 | 1.1 | | |
| $K_P(\mu\text{M})$ | 100.2 | 100.2 | 1.2 | 100.2 | | |
| $k_{cat}^F(\text{s}^{-1})$ | 2.3 | 100.3 | 1.3 | 1.3 | | |
| $k_{cat}^R(\text{s}^{-1})$ | 2.4 | 100.4 | 5.4 | 10.4 | | |
| $K_{eq}^*$ | 87 | 9.9 | 2.9 | 11.4 | | |
| $k_{cat}^F/K_S(\mu\text{M}^{-1}\text{s}^{-1})$ | 2.1 | 9.9 | 13 | 1.2 | | |

### Data set design

For the SIM1 simulations, the concentration of substrate was varied over a range of concentrations from 0.5\**K*_*S*_ to 5\**K*_*S*_ with no added product ([P_o_]) at the start of the simulations. For the SIM2 simulations, the substrate concentration was kept constant at 5\**K*_*S*_ and the concentration of initial added product at the start of each simulation was varied from 0 to 5\**K*_*P*_ (Table 2). The SIM3 simulations provided a matrix (3×3) using the variation of initial substrate ([S_o_]) and initial added product ([P_o_]) concentrations as described for SIM1 and SIM2 simulations, respectively (see Table S1 for an example of simulation conditions). The calculated time domain was optimized through the *θ* ratio of final instantaneous velocity (*v*_*f*_) to initial velocity (*v*_*o*_) to normalize the data point density across each time domain (see Materials and Methods for details). Each simulation provided a unique experimental data set with a pseudorandom Gaussian error distribution and these data sets were fit using NonlinearModelFit in *Mathematica*. For Apy and E1-E20, all four constants 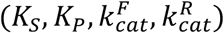 were floated during the analysis. Since Eg5 demonstrated a minimal PI mechanism, only three constants 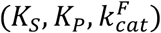 were floated during the fitting. We assessed the sensitivity of the SIM1, SIM2, and SIM3 designs against varying additive error for Apy and Eg5 (Fig 2). For Apy, SIM1 was significantly inferior to the SIM2 and SIM3 designs based on overall fitting scores (Figs 2*a–d*). However, SIM3 was a significantly better design compared to the SIM2 design (Fig 2*d*), suggesting that varying both initial substrate and product concentrations provided better overall fitting scores. For Eg5, SIM1 and SIM3 were comparable for *K*_*S*_ and 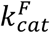 estimates (Figs 2*e*–*g*), but SIM3 provided a significantly improved mean fitting score for *K*_*P*_ (Fig 2*h*).

**FIGURE 2:**
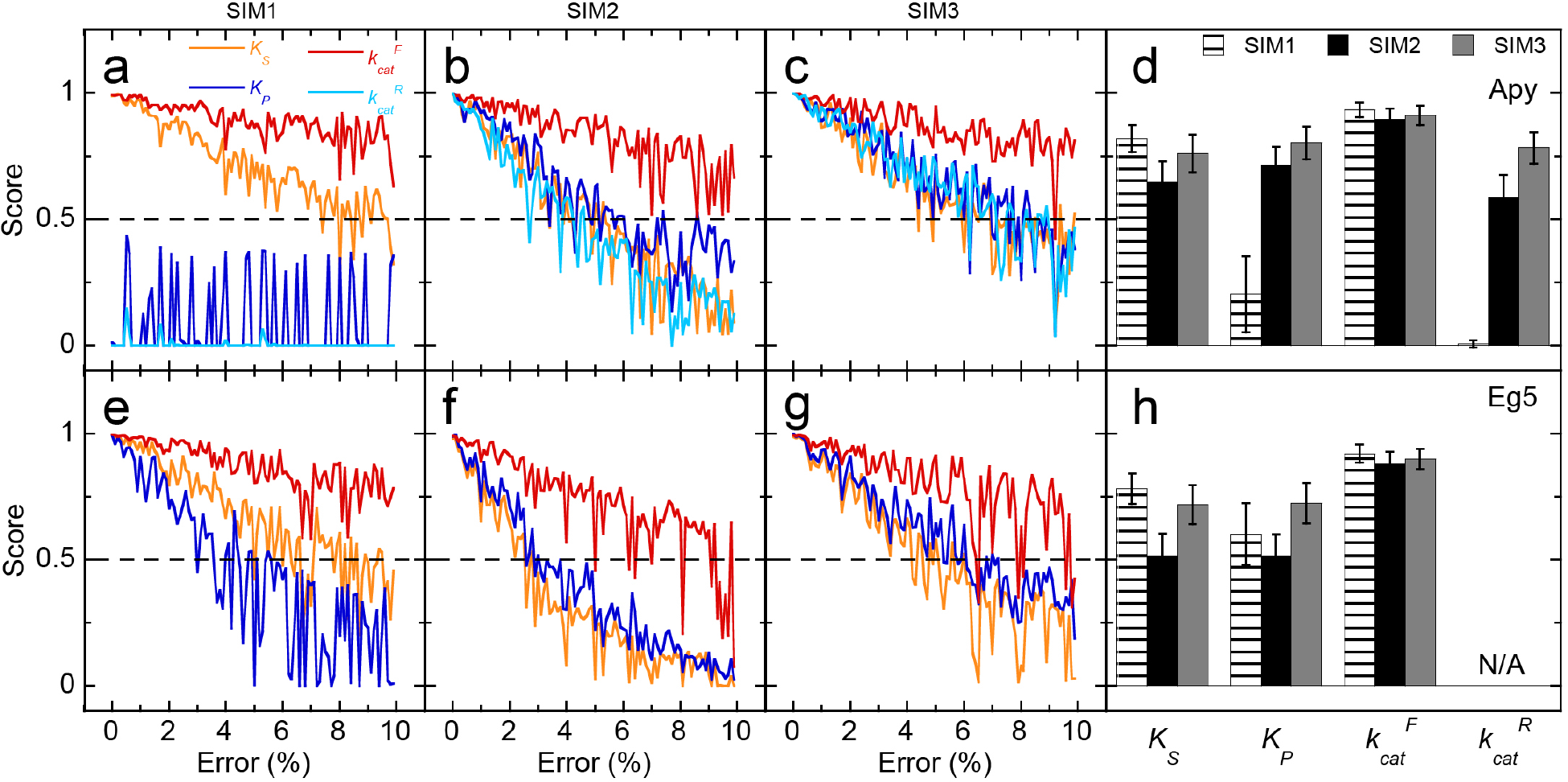
Error sensitivity for Apy and Eg5 progress curves under different simulation designs. (*a-c*) Fitting results for Apy simulations with the fitting score for each constant (*K*_*S*_, orange; *K*_*P*_, blue; *k*^*F*^_cat_, red; *k*^*R*^_cat_, cyan) plotted against error using SIM1, SIM2, and SIM3 conditions, respectively (*d*) At constant 3% error, the mean fitting score for each constant was determined under different Apy simulation designs (as indicated). n=200. Error bars represent the standard deviation of the mean. (*e-g*) Fitting results for Eg5 simulations with the fitting score for each constant plotted against error using SIM1, SIM2, and SIM3 conditions, respectively. (*h*) At constant 3% error, the mean fitting score for each constant was determined under different Eg5 simulation designs (as indicated). n=200. N/A, not applicable

We tested the sensitivity of all three simulation designs (SIM1, SIM2, SIM3) against constant additive error for our hypothetical enzymes, E1-E20 (Fig S5). For the hypothetical enzymes with *K*_*S*_ ≥ *K*_*P*_ (E1-E10), SIM2 provided very low (or indistinguishable from zero) fitting scores for *K*_*S*_ and *K*_*P*_. SIM3 provided comparable fitting scores for *K*_*S*_, *K*_*P*_ and 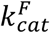, but consistently higher scores for 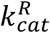. For the hypothetical enzymes with *K*_*S*_ < *K*_*P*_ (E11-E20), SIM1 provided very low (or indistinguishable from zero) fitting scores for *K*_*P*_ and 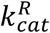. SIM3 provided consistently higher scores for all four constants compared to SIM2. There was no obvious correlation to better fitting scores when comparing 1)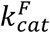 relative to 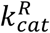, 2) overall equilibrium constant 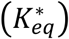, or 3) enzymatic efficiency 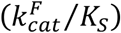. Overall, varying both initial substrate and added product was successful to attain higher fitting scores regardless of the enzyme mechanism.

We assessed the sensitivity of the SIM3 design against varying additive error for our hypothetical enzymes, E1-E20 (Fig S6). All enzymes showed fitting scores that dropped below 0.5 at >3% error except for E10, which had fitting scores that dropped below 0.5 around 2% error. Using asymmetric confidence estimate analysis, simulated Apy SIM3 designs were tested at various additive errors (Fig S7). Since the asymmetric confidence interval was consistently larger than the symmetric 95% confidence estimates obtained from nonlinear least squares fitting, it was difficult to directly compare quality scores (Fig S8). Given the wide range of different hypothetical enzymes tested in this analysis, the SIM3 design (where both initial substrate and added product is varied) is likely to be successful at estimating the four MM parameters from fairly ‘noisy’ data for any enzyme found in natural systems.

### Correlations between the estimated constants during the fitting

We next examined the six possible correlations between the four fitted parameters: 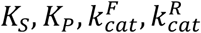 (Fig 3). Using the standard asymptotically unbiased sample correlation matrix, we computed the estimated correlations as we varied through E1-E20, Apy, and Eg5 with constant error and fixed starting values. We observed that the correlations remain roughly the same, but there was some variability as the parameters for the different enzymes vary. The relationship in the model accounts for most of the nonzero correlation especially considering a given fitting curve determined only three of the four parameters uniquely as discussed (Equations S18-S29). The simplicity of the symmetry between *K*_*S*_ and *K*_*P*_ accounts for the near perfect correlation between these two constants.

**FIGURE 3:**
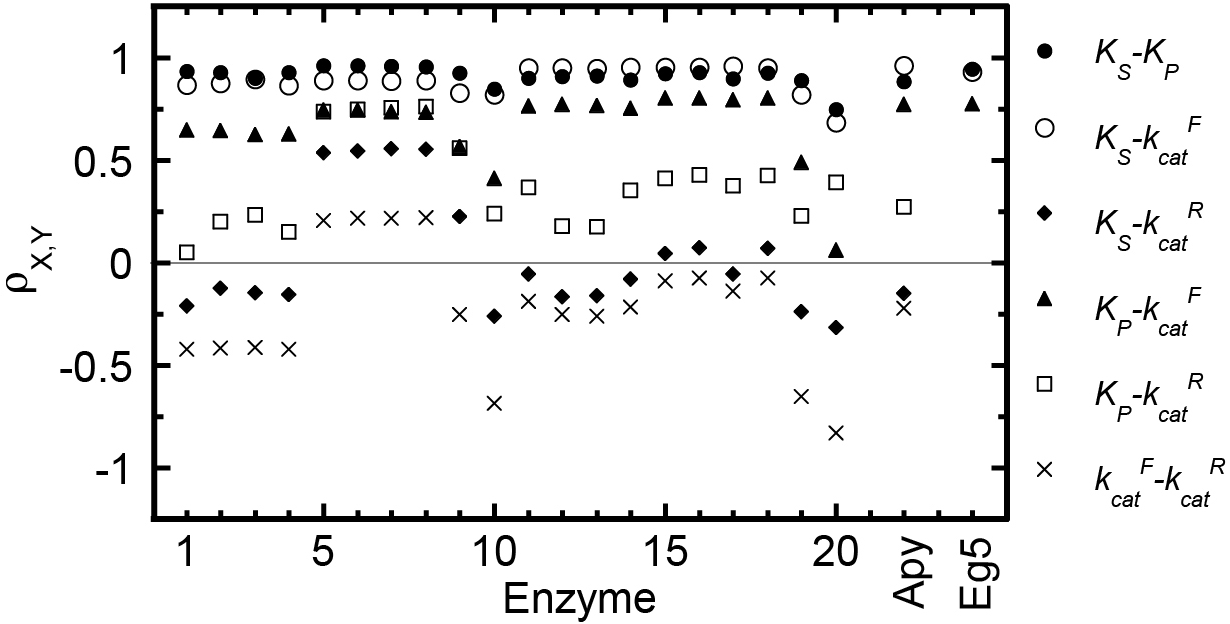
Parameter correlations for SIM3. All six parameter correlation coefficients (ρ) are shown for E1-E20, Apy, and Eg5 for the SIM3 design. Error was held constant at 3%. E1-E20 and Apy were modeled using the FR mechanism whereas Eg5 was modeled using the PI mechanism

### Time domain length effects on the parameter estimation

We investigated the sensitivity of the SIM3 design for the length of the time domain based on the ratio of instantaneous velocity from the start of the reaction till the end (*θ* = *v*_*f*_/*v*_*o*_; Fig 4). We observed similar fitting scores for Apy and Eg5 over a broad range of time domains, indicating a vast insensitivity of the SIM3 design on the short time domain for data simulation (Figs 4*a-c* and S9). At *θ* ratios greater than 0.7, the fitting scores became markedly lower, indicating that the fitting relies on a significant quantity of product formed during the reaction (i.e. 60-85% of initial substrate converted to product during the time course). However, if the time domain was extended beyond the optimal range and thus few data were found in the initial velocity phase of the curve, the fitting scores steadily decreased for Apy and Eg5 (Figs 4*d-f*). For the hypothetical enzymes with *K*_*S*_ ≥ *K*_*P*_ (E1-E10), fitting scores for *K*_*S*_ and *K*_*P*_ decreased rapidly as the time domain was lengthened (Fig S10). For the hypothetical enzymes with *K*_*S*_ < *K*_*P*_ (E11-E20), the overall fitting scores were less sensitive to the longer time domains (Fig S10).

**FIGURE 4:**
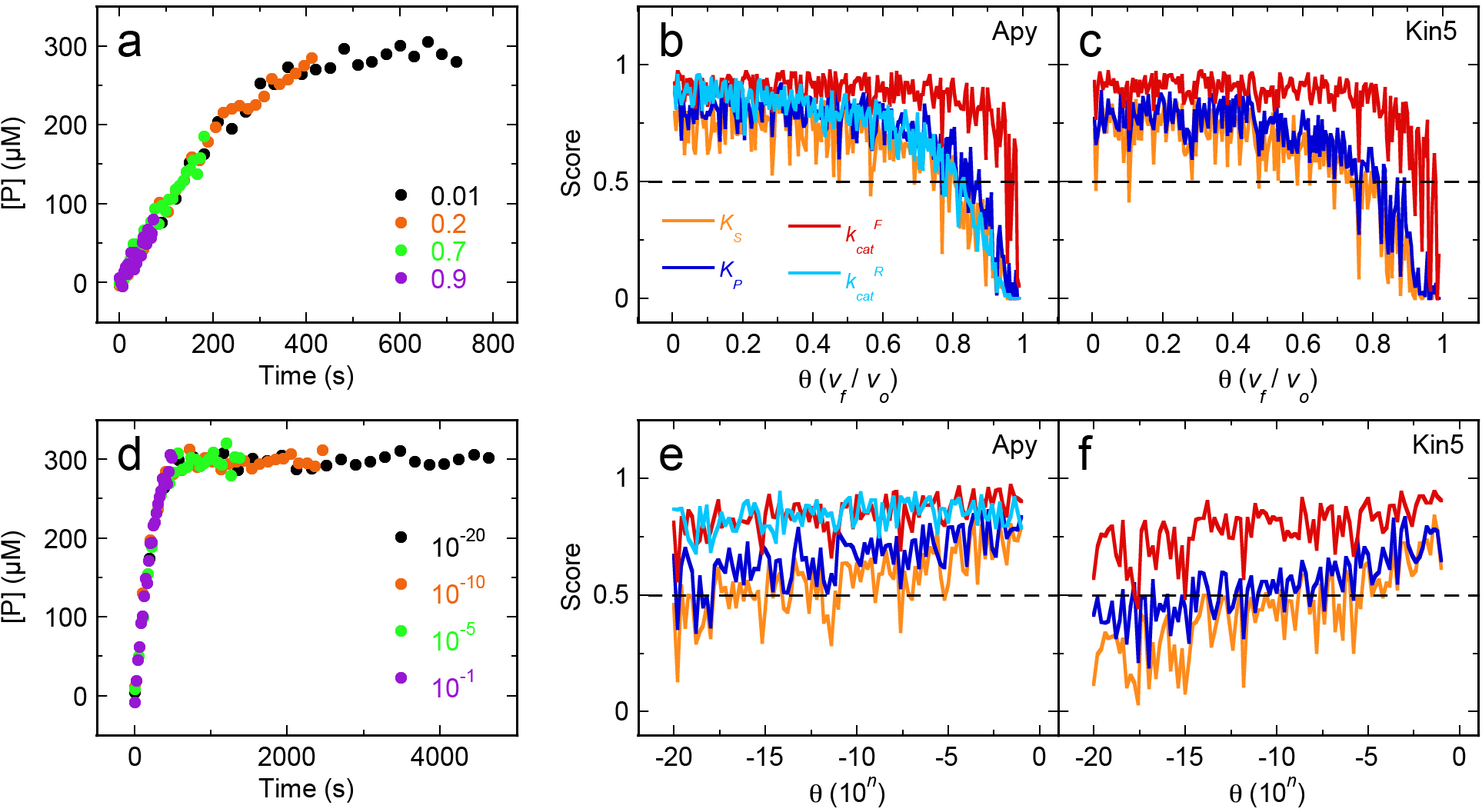
SIM3 Fitting Scores as a Function of Time Domain. Apy and Eg5 simulations were performed by varying the time domain [θ as the ratio of final instantaneous velocity (v_f_) to initial velocity (v_o_)]. (*a*) Representative Apy simulations with constant error (3%) are shown with increasing θ ratios (as indicated). (*b-c*) Fitting scores as a function of the θ ratio for Apy and Eg5, respectively, (*d*) Representative plots of Apy simulations at decreasing θ ratios, (*e-f*) Fitting cores as a function of decreasing θ values with constant error at 3% for Apy and Eg5, respectively.

### Data point density effects on parameter estimation

The total degrees of freedom correspond to the length of each progress curve (i.e. 25 data points per curve) multiplied by the number of progress curves. The number of data points per reaction was easy to modify since the total time domain was divided evenly from 2–1024 data points (i.e. changing the time constant for the calculation of [P]_*t*_). Figs 5*a-b* highlights the SIM3 results for Apy and Eg5 by varying these conditions. As expected, the fitting scores increased proportionally with the number of data points per time course. These results were not specific to Apy and Eg5, but similar observations were made for all hypothetical enzymes tested (Fig S11). We observed a similar increase of the fitting scores by holding the number of data points per curve constant at 25 and varying the number of progress curves (data not shown). However, using methods to solve the asymmetric confidence estimates, we observed nearly constant fitting score as a function of data points per curve (Fig 5*c*). Therefore, the asymmetric confidence estimates solved from fitting two constants while fixing the values of the remaining two constants provided a robust estimate of the confidence interval independent of the number of total data points in the data set.

**FIGURE 5:**
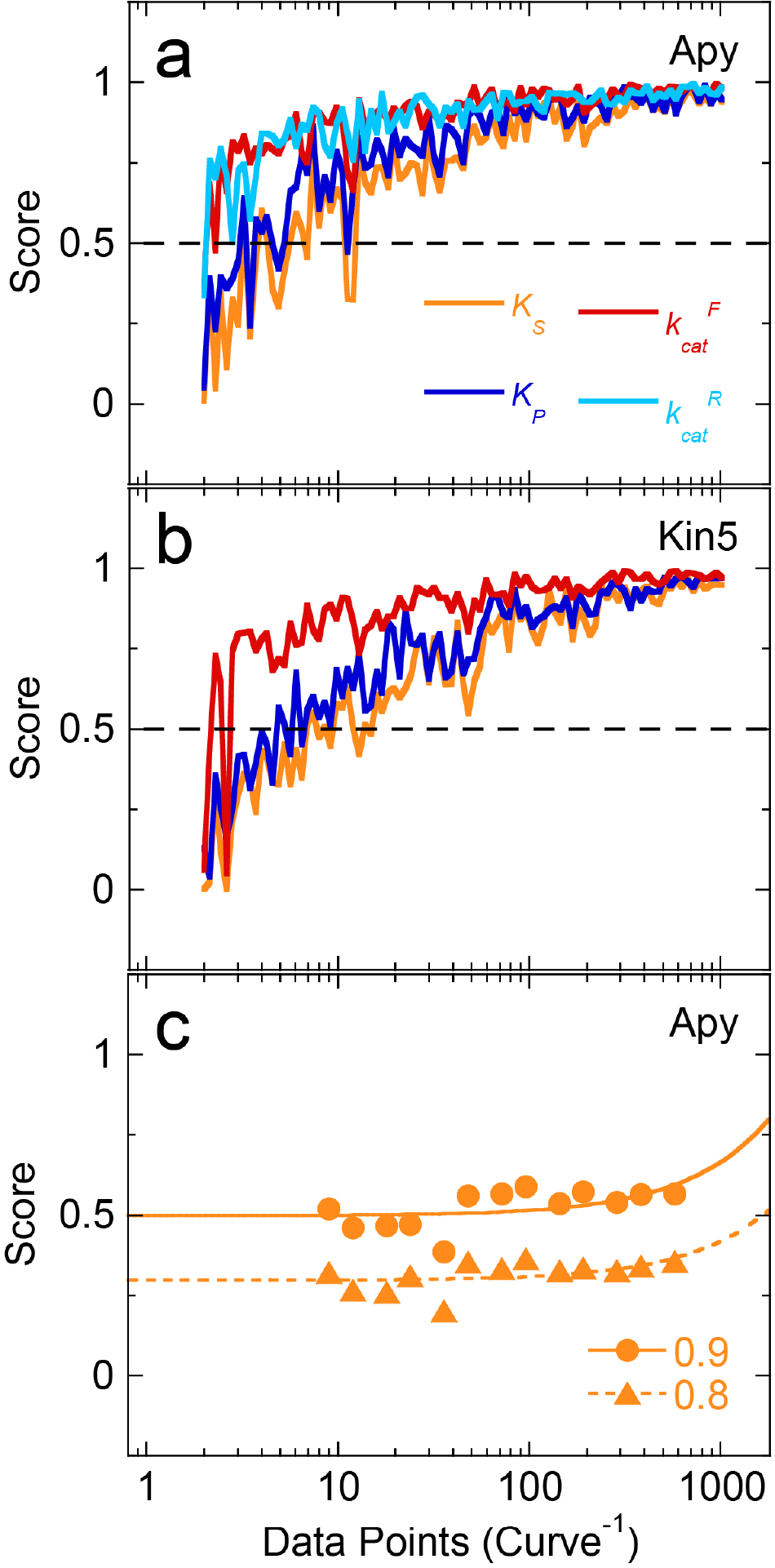
SIM3 results as a function of data point density. (*a*) Apy and (*b*) Eg5 simulations were performed by increasing the number of data points per curve (2-1024 points). Fitting scores were plotted as a function of number of data points per curve, (*c*) Asymmetric confidence estimates for *K*_*S*_ in Apy simulations at 0.8 (triangles) and 0.9 (circles) contours. Data plotted with log scale x-axis and were fit to a linear function. Error was held constant at 3%.

## Discussion

### Different enzyme mechanisms

We have tested the sensitivity of three different experimental designs centered on the closed form (Scheme 1) of the expanded Michaelis-Menten (MM) mechanism for enzyme activity. To simultaneously fit the raw data from individual progress curves to attain the four MM constants 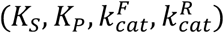, we fit real experimental data for two enzymes (Apy and Eg5) as well as simulated data for 20 different enzymes (E1–E20). These hypothetical enzymes have a wide range of equilibrium constants, substrate and product binding affinities, catalytic constants, and enzymatic efficiencies (Fig S2*a*, Table 3). Despite all the divergent mechanisms tested in this study, the SIM3 design was robust in accurately and precisely estimating all four MM constants from an experimentally feasible data set.

### Scoring function provides standard assessment

Given the complex nature of the data analysis from this study, we developed a simple exponential scoring function (Equation 9; Fig S2*b*) that yielded a score from 0 to 1 based on nonlinear least squares fitting accuracy (defined as the fit estimate divided by the actual constant) and fitting precision (defined as the symmetric 95% confidence interval for each estimate). In general, more meaningful fitted constants correlate with higher accuracy and lower confidence intervals. The weighting parameters that govern the scoring function were empirically determined to provide a fitting score of 0.5–1 when accuracy and precision were less than ±50%. The same weighting constants were used for the entire analysis, and depending on the nature of the questions being addressed, these weighting factors can be scaled accordingly to provide more stringent or more flexible scoring outcomes. The weighting factors employed in this study provide a reasonably conservative approach to evaluating the fitting results.

### Best experimental design of steady-state experiments

Our analysis took advantage of three experimental designs called SIM1 (increasing [S_o_] with no added [P_o_] at *t* = 0), SIM2 (constant [S_o_] with increasing added [P_o_] at *t* = 0), and SIM3 (matrix of increasing [S_o_] with increasing added [P_o_] at *t* = 0). Typically, steady-state experiments are designed by only varying the substrate concentration as in SIM1. However, we found that adding initial product in the beginning of the reaction will lead to better parameter fitting results from both experimental and simulated kinetic data. Indeed, the best fitting regime happened with varying both substrate and initial product concentrations in the 3×3 matrix (Fig 2). We found that >50% of initial substrate conversion to product was necessary for accurately defining all four parameters (Fig 4). If experimental error remains constant, having greater than 24 data points per curve (assuming a 3×3 matrix) provided sufficiently accurate parameter estimates, but more data narrowed the symmetric confidence limits (Fig 5). We also observed that most hypothetical enzyme mechanisms tolerated a wide range of initial substrate and initial added product concentrations for achieving accurate parameter estimates (data not shown). Given the capabilities of *Mathematica*, future investigations aim to extend this analysis to mechanisms involving two substrates and products during the reaction (i.e. bi-bi mechanisms).

### Variable conditions in the design of steady-state experiments

When studying a new uncharacterized enzyme, the decisions made by the enzymologist for initial substrate concentrations, total enzyme concentration, length of time domains, the number of data points per time domain, and the number of data sets to collect are difficult. Interestingly, global fitting under different simulation designs tolerated a wide spectrum of initial substrate and/or added product concentrations (~2-orders of magnitude around the *K*_*S*_ and *K*_*P*_ values; data not shown). The total enzyme concentration and the length of the time domain are directly related, though limits are often encountered based on the solubility limit of the enzyme, the dissociation of higher order oligomers of the enzyme, the minimum time constant for data sampling, and/or the stability of the enzyme under the experimental conditions. Therefore, these caveats make it difficult to achieve the optimal conditions to meet MM steady-state assumptions. Nevertheless, we found that our SIM3 design converged on all four parameters for all enzymes tested despite the length of time domain used.

As expected, we observed higher fitting scores when data sampling frequency increased over the course of the progress curve (Figs 5*a-b* and S11), yet, surprisingly, the scores did not linearly improve with the added number of data sets in the matrix (data not shown). These results indicate that an enzymologist can attain accurate and precise information about the kinetic mechanism with ~10-fold less time and materials. We also found the matrix of initial substrate and added product concentrations need not be symmetrical for convergence on the four parameters (data not shown). Even a minimum of 2 initial substrate concentrations and 2 added product concentrations are sufficient to attain a rough sketch of the behavior of the enzyme, which will rapidly guide the researcher to subsequent experimental conditions for the next round of experiments. This approach will provide rapid feedback to appropriately hone on the experimental conditions that provide the lowest range of confidence intervals and (hopefully) more accurate estimates of the steady-state parameters.

## Conclusions

Results presented here will impact how steady-state experiments will be designed and how steady-state kinetic data will be analyzed in the future. In the past, MM steady-state experiments focused on capturing data during the initial velocity time regime with variant substrate concentrations. These data were subsequently fit to a linear function and the slopes that define the initial reaction velocities were plotted against the substrate concentration to be fit to the hyperbolic MM equation to extract the *K*_*cat*_ *and K*_*S*_ parameters. Due to restrictions that were directly related to the experimental methodologies (e.g. linear range for product detection) or to the nature of the enzyme under investigation (e.g. an enzyme that binds its substrate very tightly but has a very slow maximal rate of substrate turnover), sometimes these experiments did not accurately define the intrinsic rate and equilibrium constants. Also, when these experiments did work as intended, the researcher would only yield two of the multiple parameters that govern the enzymatic cycle. Therefore, to minimize the time and materials used in the laboratory as well as maximize mechanistic insight through the multidimensional chemical kinetic analysis, the development of a modified experimental methodology is required. Combining the variation of both initial substrate and added product concentrations to establish a matrix of reaction conditions provides a simple strategy to generate experimental data that can be fit to extract all four MM parameters from a relatively simple data set. Since results from these experiments are typically the first insight into the mechanistic character of an unknown enzyme, having a more thorough analysis gives the enzymologist a better understanding for designing more complicated kinetic and/or thermodynamic experiments. Although these results promote the advancement of steady-state kinetic methodologies, data from presteady-state experiments are indispensible. These advancements in steady-state methodologies can provide insights of the mechanistic details that are derived through presteady-state methodologies and *vice versa*. Given the ability of *Mathematica* to fit data with pseudo-infinite dimensionality, these strategies can be applied to more complex mechanisms with multiple different substrates and products, activators, and small molecule inhibitors. Also, the progress curve fitting allows for enzyme concentration being a variable parameter, thus eliminating the need to normalize all reactions to a common enzyme concentration for comparison purposes. This study sets a foundation for modeling more complex mechanisms and will provide a new strategy for obtaining accurate and precise measurements of basic MM parameters.

## Acknowledgments

We thank Jeffrey Ewer for preliminary data collection as well as Kayla Bell and Benjamin Walker for insightful comments on the manuscript. We acknowledge financial support from the Simons Foundation (grant #210442 to C.G.C.), the IU Faculty Research Support Program-External Resubmission (J.C.C.) and the National Science Foundation (MCB 1614514 to J.C.C.).

## Abbreviations

^1^ MM: Michaelis-Menten
E: enzyme
S: substrate
P: product
errA: additive error
P_i_: inorganic phosphate
*C*_*est*_: estimated constant from nonlinear least squares fitting
*C*_*exp*_: the expected constant
*v*_*f*_: final instantaneous velocity
*v*_*o*_: initial velocity
CI: confidence interval (95%)
AIC: Akaike information criterion
BIC: Bayesian information criterion
MT: microtubule.

