## Supplemental Info for "Error sensitivity and optimization of steady-state kinetic parameters using multidimensional chemical kinetic analysis"

### Supporting information

#### Derivation of the closed form of the model

Michaelis and Menten proposed a simple two-step mechanism (Equation 1) to explain the kinetic behavior of the enzyme invertase with reversible substrate binding and irreversible chemistry coupled to product release [1]. Countless investigators have utilized this mechanism over the past century, yet the simple model does not consider competitive product inhibition of the enzyme activity (Equation 2) and/or reversible chemistry (Equation 3). Based on Equation 3, the total enzyme ($\mathrm{E_o}$) can be described as a function of different bound states as follows:

(S1)

$$[\mathrm{E_o}] = [\mathrm{E}]_{\mathrm{free}} + [\mathrm{ES}] + [\mathrm{EP}]$$

According to the steady-state assumption, the time derivatives of each enzyme complex can be set equal to zero:

(S2)

$$\frac{d[\mathrm{ES}]}{dt} = k_1[\mathrm{S}][\mathrm{E}]_{\mathrm{free}} + k_4[\mathrm{EP}] - (k_2 + k_3)[\mathrm{ES}]$$

(S3)

$$\frac{d[\mathrm{EP}]}{dt} = k_6[\mathrm{P}][\mathrm{E}]_{\mathrm{free}} + k_3[\mathrm{ES}] - (k_4 + k_5)[\mathrm{EP}]$$

Solving Equations S1–S3 for the [S] and [P] yields the subsequent fully reversible MM rate law:

(S4)

$$v(t) = [\mathrm{E_o}]\frac{k_{cat}^{F}\frac{[\mathrm{S}]}{K_S} - k_{cat}^{R}\frac{[\mathrm{P}]}{K_P}}{1 + \frac{[\mathrm{S}]}{K_S} + \frac{[\mathrm{P}]}{K_P}}$$

where $K_S$ and $K_P$ are the equilibrium binding constants for substrate and product, and $k_{cat}^{F}$ and $k_{cat}^{R}$ are the forward and reverse catalytic constants, respectively. According to the rate law provided in Equation S4, these four constants can be arranged into the Haldane relation: a basic thermodynamic constant representing the ratio between substrate and product concentrations at equilibrium:

(S5)

$$K_{eq}^{*} = \frac{k_{cat}^{F}K_P}{k_{cat}^{R}K_S}$$

We can rearrange Equation S4 and substitute Equation S5 to describe the rate law for a competitive product inhibition model [2]:

(S6)

$$\frac{d[\mathrm{P}]}{dt} = \frac{k_{cat}^{F}\,[\mathrm{E_o}]\left[[\mathrm{S}] - \frac{[\mathrm{P}]}{K_{eq}^{*}}\right]}{K_S\left(1 + \frac{[\mathrm{S}]}{K_S} + \frac{[\mathrm{P}]}{K_P}\right)}$$

The relationship between the substrate and product concentrations over time relates to the following equations:

$$[\mathrm{S}]_t = [\mathrm{S_o}] - [\mathrm{P}]_t \tag{S7}$$

$$[\mathrm{P}]_t^* = [\mathrm{P}]_t + [\mathrm{P_o}] \tag{S8}$$

where $[\mathrm{S}]_t$ and $[\mathrm{P}]_t$ are the substrate and product concentrations (respectively) at time *t*, $[\mathrm{S_o}]$ and $[\mathrm{P_o}]$ are the initial substrate and added product concentrations (respectively) at time = 0, and $[\mathrm{P}]_t^*$ is the total product at time *t*.

Using *Mathematica* 10 (Wolfram), it was verified by differentiation in time that the differential equation solution takes the following closed form, where $\omega$ represents the Lambert omega-function [3-12].

$$\begin{aligned}[\mathrm{P}]_t &= \left(\frac{1}{(K_P - K_S)(k_{cat}^F K_P + k_{cat}^R K_S)}\right) \Bigg\{ (K_P - K_S)([\mathrm{S_o}]k_{cat}^F K_P - [\mathrm{P_o}]k_{cat}^R K_S) \\ &- K_P K_S([\mathrm{P_o}](k_{cat}^R + k_{cat}^F) + [\mathrm{S_o}](k_{cat}^R + k_{cat}^F) + k_{cat}^F K_P + k_{cat}^R K_S) \\ &* \omega\left[\frac{e^{-\frac{(K_P - K_S)(-[\mathrm{S_o}]k_{cat}^F K_P + [\mathrm{P_o}]k_{cat}^R K_S) + [\mathrm{E_o}](k_{cat}^F K_P + k_{cat}^R K_S)^2 t}{K_P K_S([\mathrm{P_o}](k_{cat}^R + k_{cat}^F) + [\mathrm{S_o}](k_{cat}^R + k_{cat}^F) + k_{cat}^F K_P + k_{cat}^R K_S)}} (K_P - K_S)([\mathrm{S_o}]k_{cat}^F K_P - [\mathrm{P_o}]k_{cat}^R K_S)}{K_P K_S([\mathrm{P_o}](k_{cat}^R + k_{cat}^F) + [\mathrm{S_o}](k_{cat}^R + k_{cat}^F) + k_{cat}^F K_P + k_{cat}^R K_S)}\right]\Bigg\}\end{aligned} \tag{S9}$$

In fact, the Lambert omega-function is the inverse function to $ze^z$ and since this function is two to one, its inverse has two branches $\omega_0$ and $\omega_{-1}$, which are characterized by being at least -1 or at most -1, respectively. In our case, we are using $\omega = \omega_0$ in the solution above. This can be verified by evaluating at $t = 0$ whence in Equation S9 using our convention,

$$\begin{aligned}&\omega\left[\frac{e^{-\frac{(K_P - K_S)(-[\mathrm{S_o}]k_{cat}^F K_P + [\mathrm{P_o}]k_{cat}^R K_S)}{K_P K_S([\mathrm{P_o}](k_{cat}^R + k_{cat}^F) + [\mathrm{S_o}](k_{cat}^R + k_{cat}^F) + k_{cat}^F K_P + k_{cat}^R K_S)}} (K_P - K_S)([\mathrm{S_o}]k_{cat}^F K_P - [\mathrm{P_o}]k_{cat}^R K_S)}{K_P K_S([\mathrm{P_o}](k_{cat}^R + k_{cat}^F) + [\mathrm{S_o}](k_{cat}^R + k_{cat}^F) + k_{cat}^F K_P + k_{cat}^R K_S)}\right] \\ &= \frac{(K_P - K_S)([\mathrm{S_o}]k_{cat}^F K_P - [\mathrm{P_o}]k_{cat}^R K_S)}{K_P K_S([\mathrm{P_o}](k_{cat}^R + k_{cat}^F) + [\mathrm{S_o}](k_{cat}^R + k_{cat}^F) + k_{cat}^F K_P + k_{cat}^R K_S)} \geq -1\end{aligned} \tag{S10}$$

The last inequality holds since after multiplying through by the denominator and canceling terms we arrive at $(k_{cat}^F K_P + k_{cat}^R K_S)(K_S[\mathrm{P_o}] + K_P(K_S + [\mathrm{S_o}])) \geq 0$ which holds since all terms on the left hand side are nonnegative. This implies $[\mathrm{P}]_0 = 0$ as desired. Had we incorrectly chosen the branch $\omega = \omega_{-1}$ then the value of $\omega$ would be less than -1 resulting in a nonzero value for $[\mathrm{P}]_0$. Note however, that there are closely related MM models where the $\omega_{-1}$ branch appears as the solution (e.g. see [43]), however either the underlying differential equation or the assumptions on parameters and initial conditions differ in those cases.

Here we list the relationship between the various constants that arise in the derivation of the MM model as well as the closed form of the model itself. Constants derived from mass-action kinetic parameters of Equation 3.

$$K_S = \frac{k_2k_4 + k_2k_5 + k_3k_5}{k_1(k_3 + k_4 + k_5)} \tag{S11}$$

$$K_P = \frac{k_2k_4 + k_2k_5 + k_3k_5}{k_6(k_2 + k_3 + k_4)} \tag{S12}$$

$$k_{cat}^F = \frac{k_3k_5}{k_3 + k_4 + k_5} \tag{S13}$$

$$k_{cat}^R = \frac{k_2k_4}{k_2 + k_3 + k_4} \tag{S14}$$

Solving individual Haldane constants derived from mass-action kinetic parameters of Equation 3 using Equation S9.

$$k_2 = \frac{k_1K_S(k_{cat}^R + k_6K_P) - k_{cat}^F k_6K_P}{k_6K_P - k_{cat}^R - k_{cat}^F} \tag{S15}$$

$$k_3 = \frac{k_{cat}^F k_6K_P}{k_6K_P - k_{cat}^R - k_{cat}^F} \tag{S16}$$

$$k_4 = \frac{k_{cat}^R k_1K_S}{k_1K_S - k_{cat}^R - k_{cat}^F} \tag{S17}$$

$$k_5 = \frac{k_1K_S(k_{cat}^R + k_6K_P) - k_{cat}^F k_6K_P}{k_1K_S - k_{cat}^R - k_{cat}^F} \tag{S18}$$

### Parameter Redundancy

During early attempts to accurately fit the MM model we noticed a wide range of parameter dependency yielding very similar curves. We soon discovered an intrinsic hidden parameter redundancy that occurs in the closed form solution of Equation S9 among the four constants $K_S, K_P, k_{cat}^F, k_{cat}^R$ and the initial substrate and product $[\mathrm{S_o}], [\mathrm{P_o}]$ and enzyme $[\mathrm{E_o}]$. Fortunately, this redundancy can also be expressed in closed form, and we were able to establish that this is the unique internal redundancy. Namely, while the profile curve $[\mathrm{P}]_t$ is uniquely determined by the three quantities α, β and γ (Scheme 1), there is an internal redundancy among these when we express everything in terms of the rate constants. More specifically, given the three parameters α, β, and γ we can keep them constant while varying the rate constants if and only if they are related in precisely the following way:

$$K_S = \frac{K_P\,(\alpha + \beta - [\mathrm{S_o}])}{[\mathrm{P_o}] + K_P + \alpha + \beta} \tag{S19}$$

$$k_{cat}^{F} = \frac{\gamma([\mathrm{P_o}] + [\mathrm{S_o}] + K_P)([\mathrm{P_o}] + \alpha)}{[\mathrm{E_o}]\,([\mathrm{P_o}] + [\mathrm{S_o}])([\mathrm{P_o}] + K_P + \alpha + \beta)} \tag{S20}$$

$$k_{cat}^{R} = \frac{\gamma([\mathrm{P_o}] + [\mathrm{S_o}] + K_P)\,([\mathrm{S_o}] - \alpha)}{[\mathrm{E_o}]\,([\mathrm{P_o}] + [\mathrm{S_o}])(\alpha + \beta - [\mathrm{S_o}])} \tag{S21}$$

If we want to further assume that the rate constants and initial concentrations are nonnegative, then we must further require the restrictions: $\gamma \geq 0$ and $\alpha + \beta \geq [\mathrm{S_o}]$ or $\gamma < 0$ and $\alpha + \beta < -[\mathrm{P_o}] - K_P$ as well as $[\mathrm{S_o}] \geq \alpha \geq -[\mathrm{P_o}]$. The derivation of all the above follows from simply reducing the three equations for the constants α, β, and γ and eliminating parameters. We can then easily check for the sign parameters, which are cyclic in these equations. In particular, we may treat $K_P$ as a free parameter in these equations. Specifying $K_P$ will lock the other three constant for a given curve and choice of initial quantities for product, substrate and enzyme. This is critical for understanding how to apply this to experimental design. Since these relations only involve fractional linear transformations, they can be rewritten so that any of the other variables becomes the free parameter.

For $K_S$ as the free parameter we have:

$$K_P = \frac{K_S(\alpha + \beta + [\mathrm{P_o}])}{\alpha + \beta - K_S - [\mathrm{S_o}]} \tag{S22}$$

$$k_{cat}^{F} = \frac{\gamma(\alpha + [\mathrm{P_o}])(K_S + [\mathrm{P_o}] + [\mathrm{S_o}])}{[\mathrm{E_o}](\alpha + \beta + [\mathrm{P_o}])([\mathrm{P_o}] + [\mathrm{S_o}])} \tag{S23}$$

$$k_{cat}^{R} = \frac{\gamma([\mathrm{S_o}] - \alpha)(K_S + [\mathrm{P_o}] + [\mathrm{S_o}])}{[\mathrm{E_o}](\alpha + \beta - K_S - [\mathrm{S_o}])([\mathrm{P_o}] + [\mathrm{S_o}])} \tag{S24}$$

For $k_{cat}^{F}$ as the free parameter we have:

$$K_P = \frac{([\mathrm{E_o}]k_{cat}^{F}(\alpha + \beta + [\mathrm{P_o}]) - \gamma(\alpha + [\mathrm{P_o}]))([\mathrm{P_o}] + [\mathrm{S_o}])}{\gamma(\alpha + [\mathrm{P_o}]) - [\mathrm{E_o}]k_{cat}^{F}([\mathrm{P_o}] + [\mathrm{S_o}])} \tag{S25}$$

$$K_S = \frac{([\mathrm{E_o}]k_{cat}^{F}(\alpha + \beta + [\mathrm{P_o}]) - \gamma(\alpha + [\mathrm{P_o}]))([\mathrm{P_o}] + [\mathrm{S_o}])}{\gamma(\alpha + [\mathrm{P_o}])} \tag{S26}$$

$$k_{cat}^{R} = \frac{\gamma k_{cat}^{F}([\mathrm{S_o}] - \alpha)}{\gamma(\alpha + [\mathrm{P_o}]) - [\mathrm{E_o}]k_{cat}^{F}([\mathrm{P_o}] + [\mathrm{S_o}])} \tag{S27}$$

For $k_{cat}^R$ as the free parameter we have:

$$K_P = \frac{([P_o] + [S_o])(\gamma([S_o] - \alpha) - [E_o]k_{cat}^R(\alpha + \beta - [S_o]))}{\gamma(\alpha - [S_o])} \tag{S28}$$

$$K_S = \frac{([E_o]k_{cat}^R(\alpha + \beta - [S_o]) - \gamma([S_o] - \alpha))([P_o] + [S_o])}{\gamma([S_o] - \alpha) + [E_o]k_{cat}^R([P_o] + [S_o])} \tag{S29}$$

$$k_{cat}^F = \frac{\gamma k_{cat}^R(\alpha + [P_o])}{\gamma([S_o] - \alpha) + [E_o]k_{cat}^R([P_o] + [S_o])} \tag{S30}$$

Among various trials conducted, we also explore how varying one of the initial concentrations along with time allows us to determine all the constants even under a sizable multiplicative and additive error. In fact, we examine recovery of constants under error conditions that should exceed any moderately controlled lab experiments. The most important point of the explicit parameter redundancy calculations is that we know precisely how to avoid unnecessary experiments that are of no further value in fitting the model because they fail to help pinpoint the position along the one-dimensional curve (resulting from one degree of freedom) where the coupled parameters reside.

Table S1. **Simulation conditions for Eg5.**

| **SIM1** | Curve 1 | Curve 2 | Curve 3 | Curve 4 | Curve 5 | Curve 6 | Curve 7 | Curve 8 | Curve 9 |
|---|---|---|---|---|---|---|---|---|---|
| $[S_o]$ | 15 μM | 32 μM | 49 μM | 66 μM | 83 μM | 99 μM | 116 | 133 | 150 μM |
| $[P_o]$ | 0 μM | 0 μM | 0 μM | 0 μM | 0 μM | 0 μM | 0 μM | 0 μM | 0 μM |

| **SIM2** | Curve 1 | Curve 2 | Curve 3 | Curve 4 | Curve 5 | Curve 6 | Curve 7 | Curve 8 | Curve 9 |
|---|---|---|---|---|---|---|---|---|---|
| $[S_o]$ | 150 μM | 150 μM | 150 μM | 150 μM | 150 μM | 150 μM | 150 μM | 150 μM | 150 μM |
| $[P_o]$ | 0 μM | 150 μM | 300 μM | 450 μM | 600 μM | 750 μM | 900 μM | 1050 μM | 1200 μM |

| **SIM3** | Curve 1 | Curve 2 | Curve 3 | Curve 4 | Curve 5 | Curve 6 | Curve 7 | Curve 8 | Curve 9 |
|---|---|---|---|---|---|---|---|---|---|
| $[S_o]$ | 15 μM | 83 μM | 150 μM | 15 μM | 83 μM | 150 μM | 15 μM | 83 μM | 150 μM |
| $[P_o]$ | 0 μM | 0 μM | 0 μM | 600 μM | 600 μM | 600 μM | 1200 μM | 1200 μM | 1200 μM |

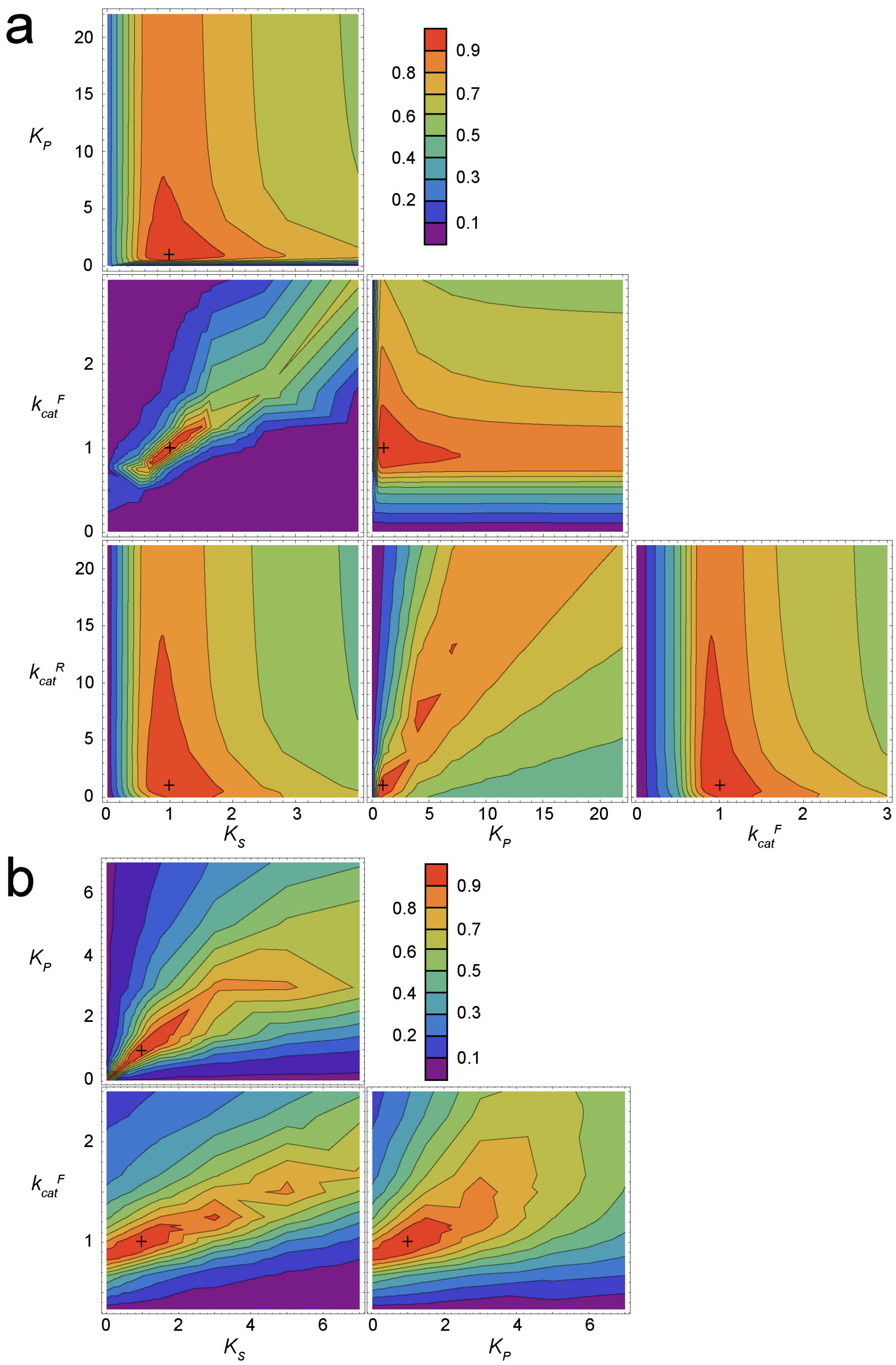


FIGURE S1: **Asymmetric confidence estimates for Apy and Eg5.** Contour plots showing $\text{MSE}_{min}/\text{MSE}_{x,y}$ for (*a*) Apy and (*b*) Eg5 experimental data presented in Fig 1 with two fixed constants and two floated constants. Axes correspond to $C_{fixed}/C_{best}$ with crosses at corresponding pairs of best fit constants.

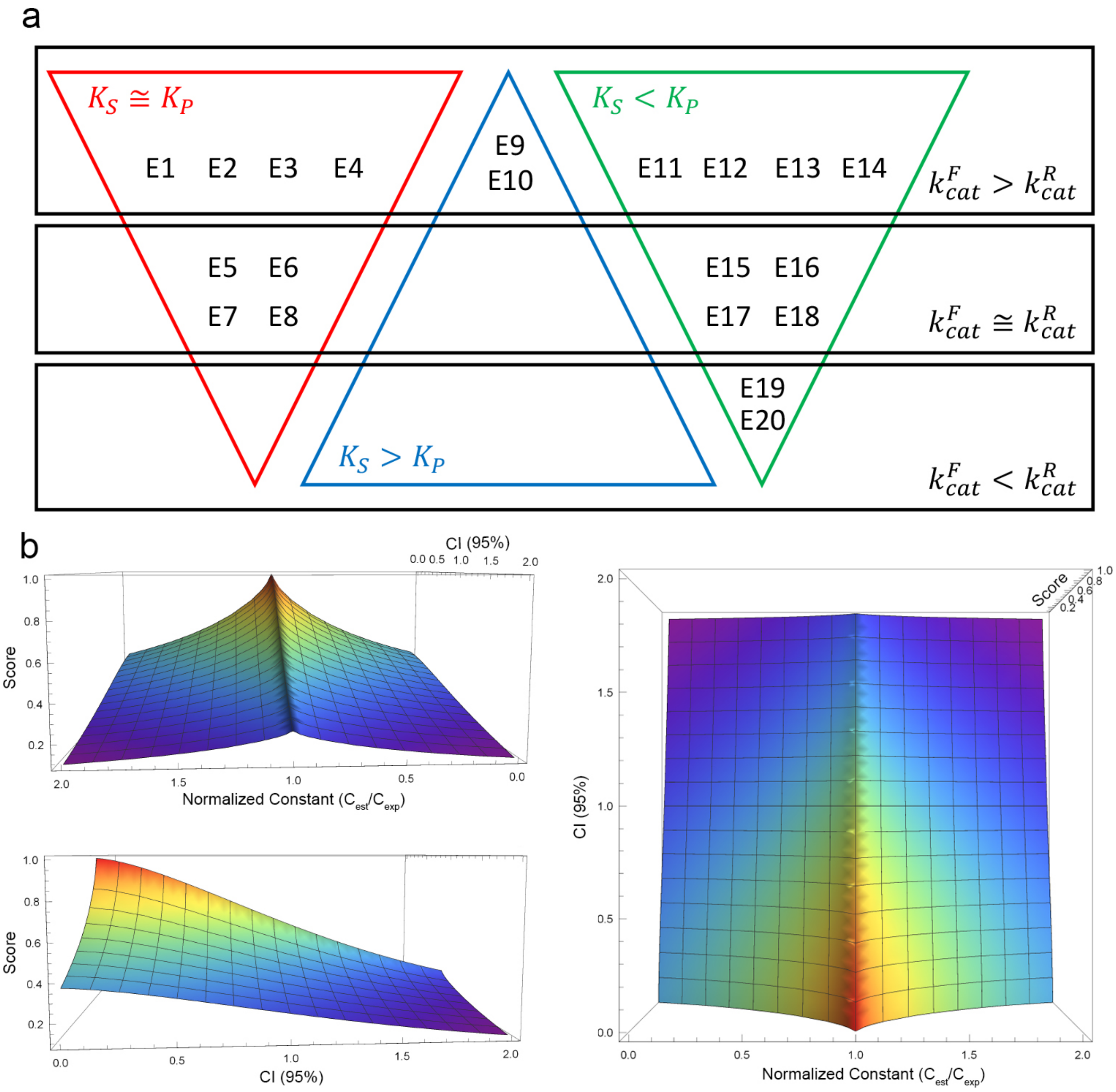


FIGURE S2: **Hypothetical enzymes and scoring function for nonlinear least squares fitting.** (*a*) Venn diagram for the tested hypothetical enzymes. (*b*) Three different views of the three-dimensional plot of the fitting score versus the normalized constant ($C_{est}/C_{exp}$) and confidence interval (95%). The surface of the plot is rainbow colored per score magnitude (i.e. red to violet for high to low scores, respectively).

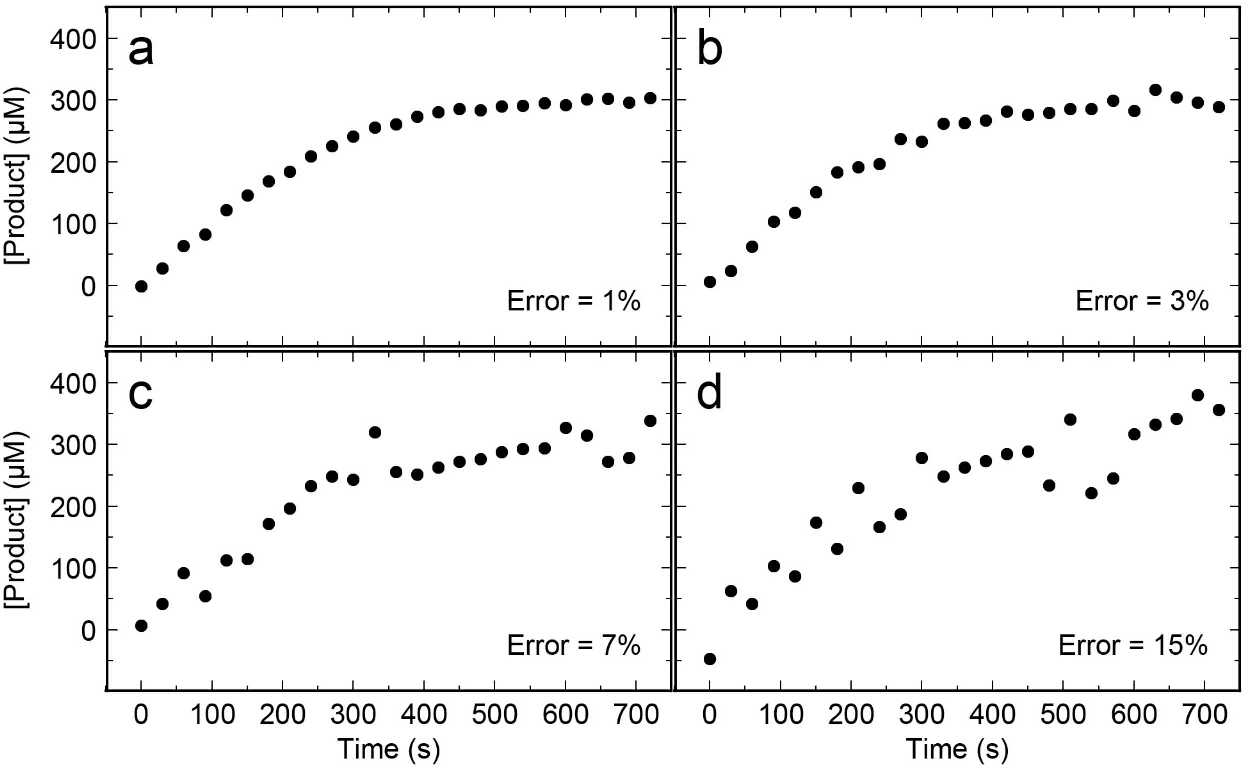


FIGURE S3: **Representative simulated data versus error.** Plots of simulated Apy product formation as a function of time are shown at different additive errors (as indicated in each panel). Each progress curve contains 25 data points. Simulation conditions: 0.08 μM Apy, 300 μM [$S_o$], 0 μM [$P_o$]. Constants found in Table 3.

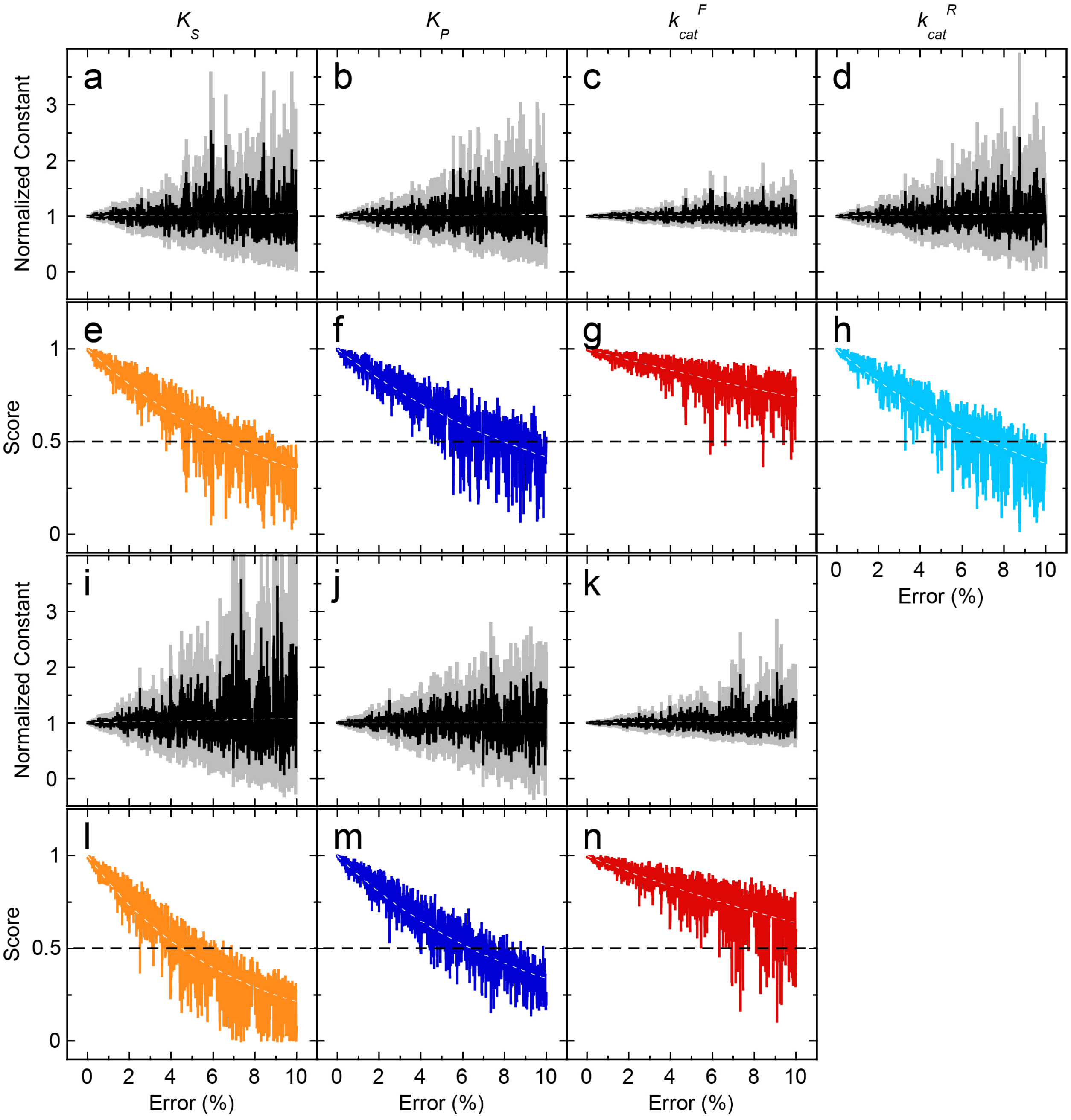


FIGURE S4: **Relationship between accuracy and precision of fitting relative to the scoring function.** (*a-d*) Results from a representative Apy simulation as a function of error using the FR model. The black line corresponds to each normalized fitted constant ($C_{est}/C_{exp}$) with the gray line representing the symmetric 95% confidence interval. The dotted lines were the fits of the normalized constants to a linear function. (*e-h*) Fitting scores (Equation 9) were plotted against error, with the dotted lines as fits to an exponential function. (*i-n*) Results from an Eg5 simulation as a function of error using the PI model. Panels are similar to (*a-h*).

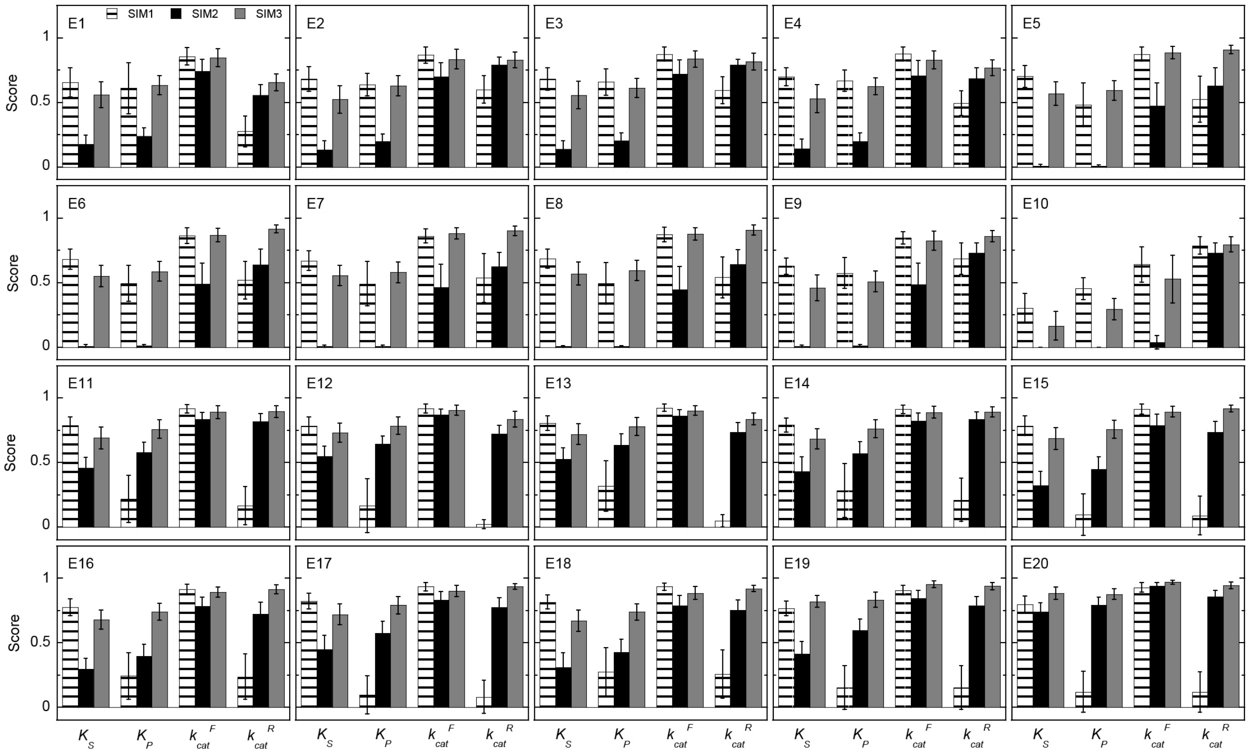


FIGURE S5: **Error sensitivity for E1-E20 progress curves under different simulation designs.** At constant 3% error, the mean fitting score for each constant was determined under different simulation designs (SIM1, SIM2, SIM3). *n*=50. Error bars represent the standard deviation of the mean score.

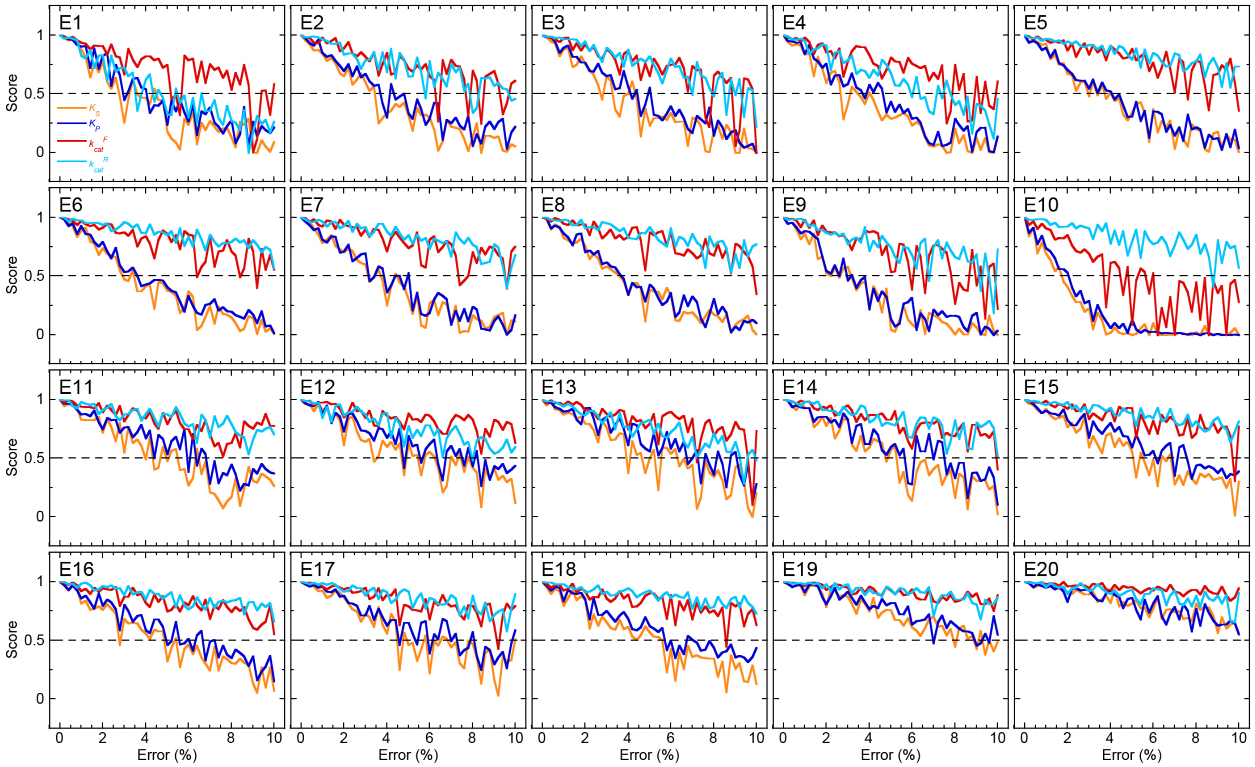


FIGURE S6: **E1-E20 error sensitivity using the SIM3 conditions.** Each panel shows the fitting scores for each constant ($K_S$, orange; $K_P$, blue; $k_{cat}^F$, red; $k_{cat}^R$, cyan) were plotted as a function of error. The x- and y-axes in each panel are scaled identically.

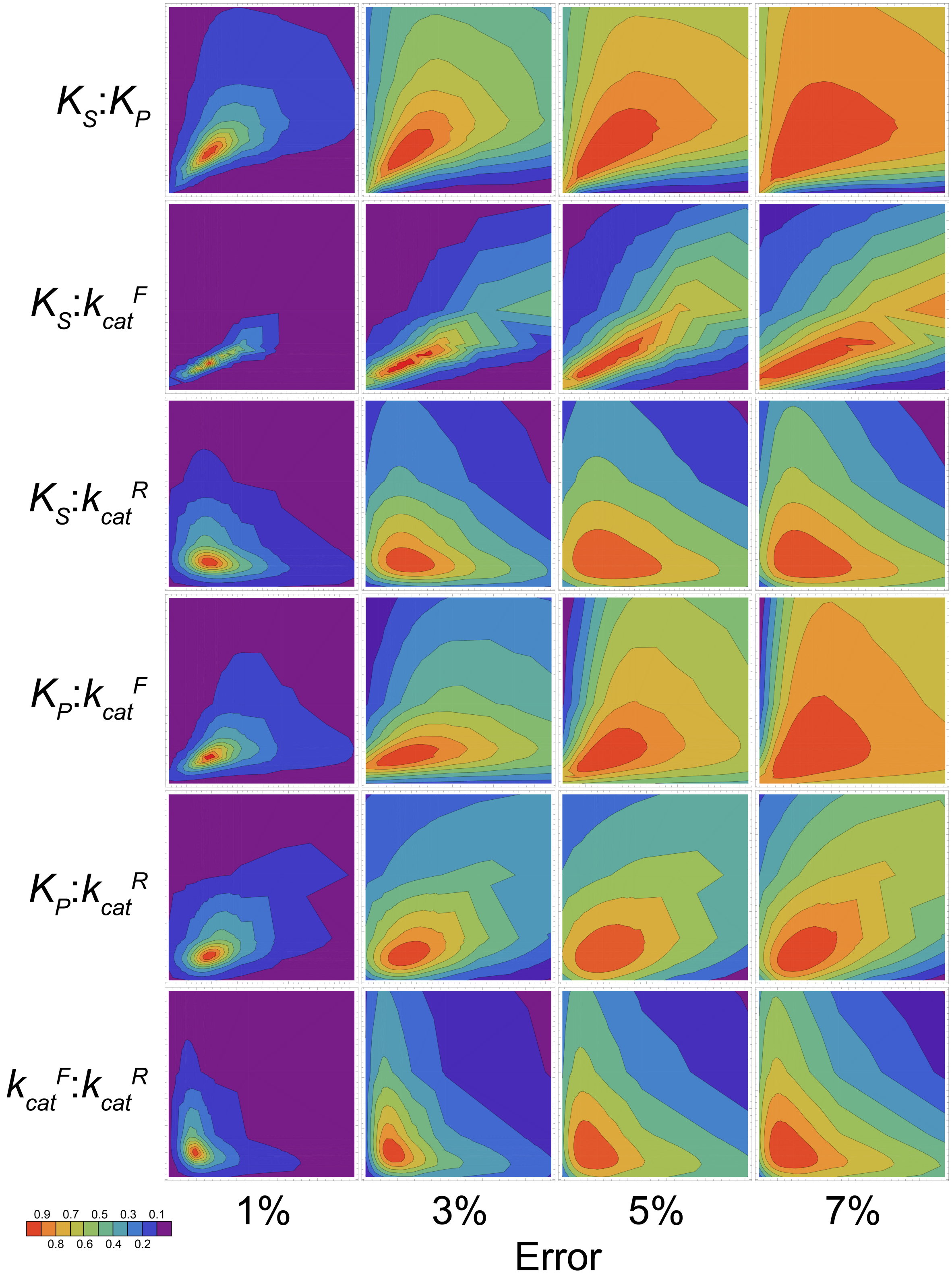


FIGURE S7: **Asymmetric confidence estimates for Apy versus error using SIM3 conditions.** Contour plots showing $MSE_{min}/MSE_{x,y}$ for Apy SIM3 data with two fixed constants and two floated constants. Axes correspond to $C_{fixed}/C_{best}$.

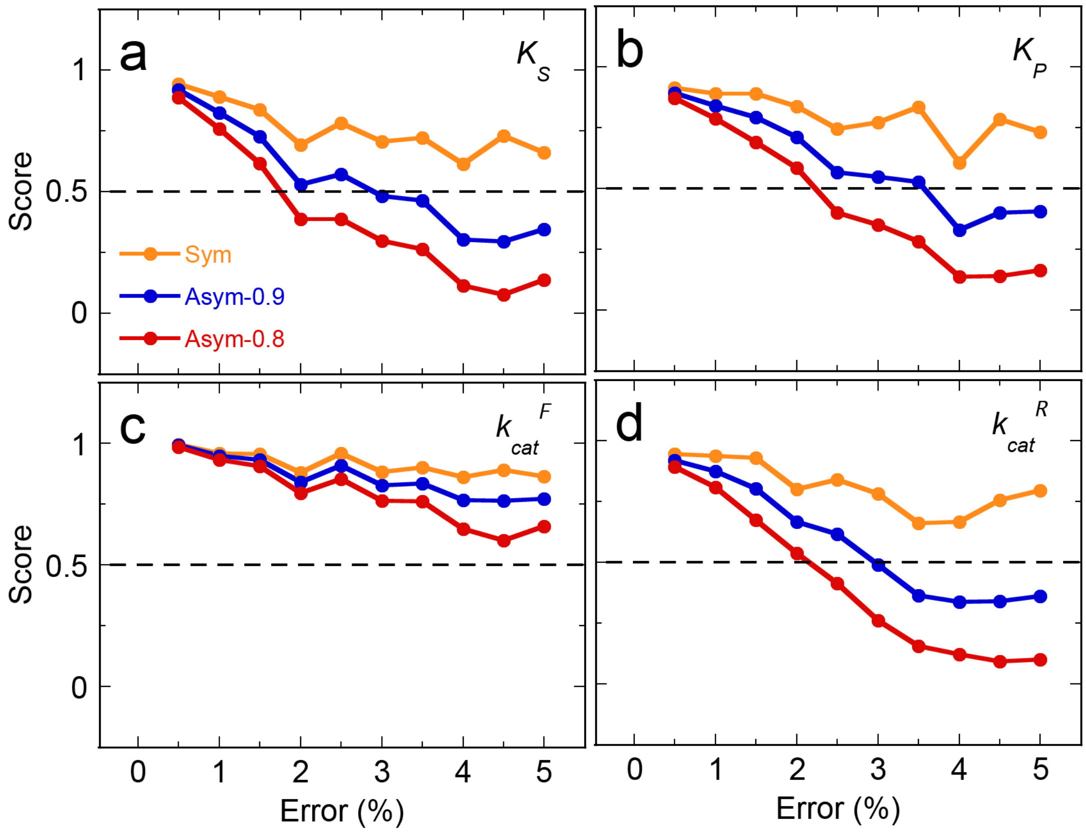


FIGURE S8: **Comparison of fitting scores based on symmetric and asymmetric confidence estimates for Apy using SIM3 conditions.** Estimates for confidence intervals for Apy SIM3 designs based on symmetric 95% CI, asymmetric 0.9 contours, and asymmetric 0.8 contours (as indicated).

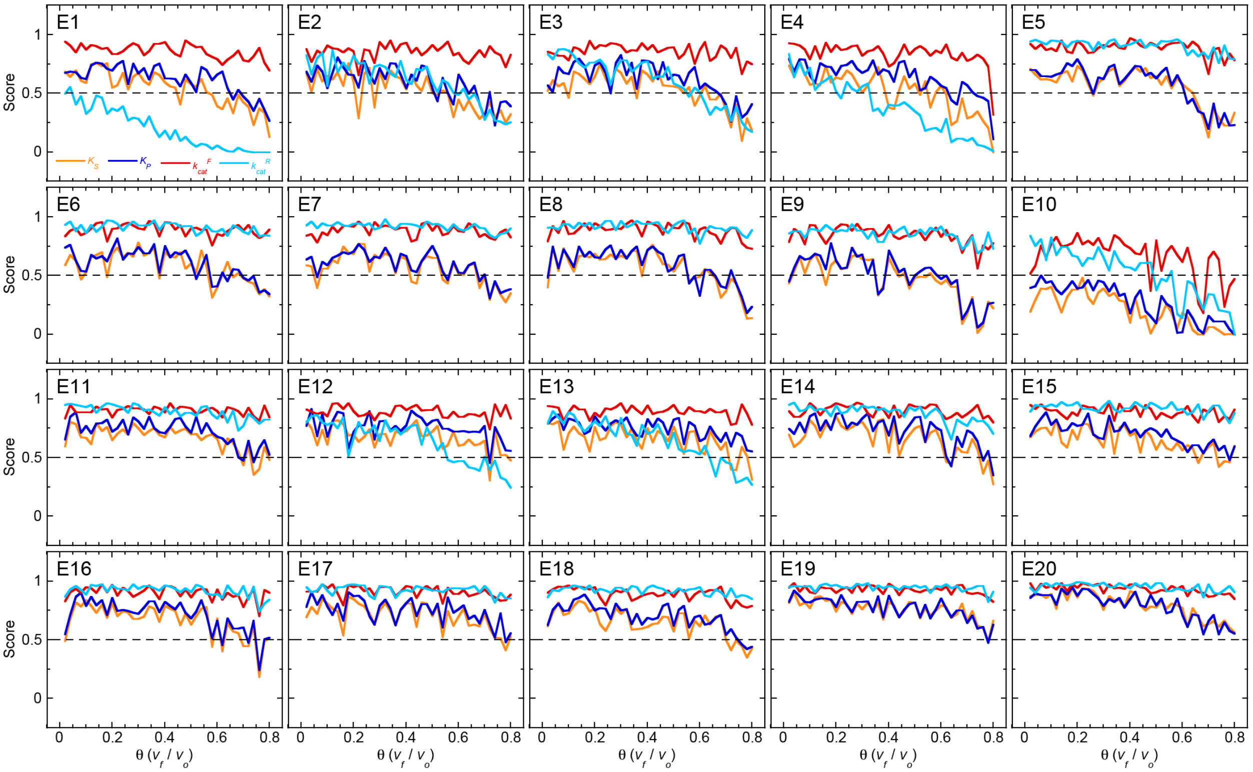


FIGURE S9: **Results for E1-E20 as a function of short simulation time domains.** E1–E20 simulations were performed by varying the time domain [$\theta$ as the ratio of final instantaneous velocity ($v_f$) to initial velocity ($v_o$)]. E1-E20 fitting scores for each constant ($K_S$, orange; $K_P$, blue; $k_{cat}^{F}$, red; $k_{cat}^{R}$, cyan) were plotted as a function of the $\theta$ ratio with error held constant at 3%.

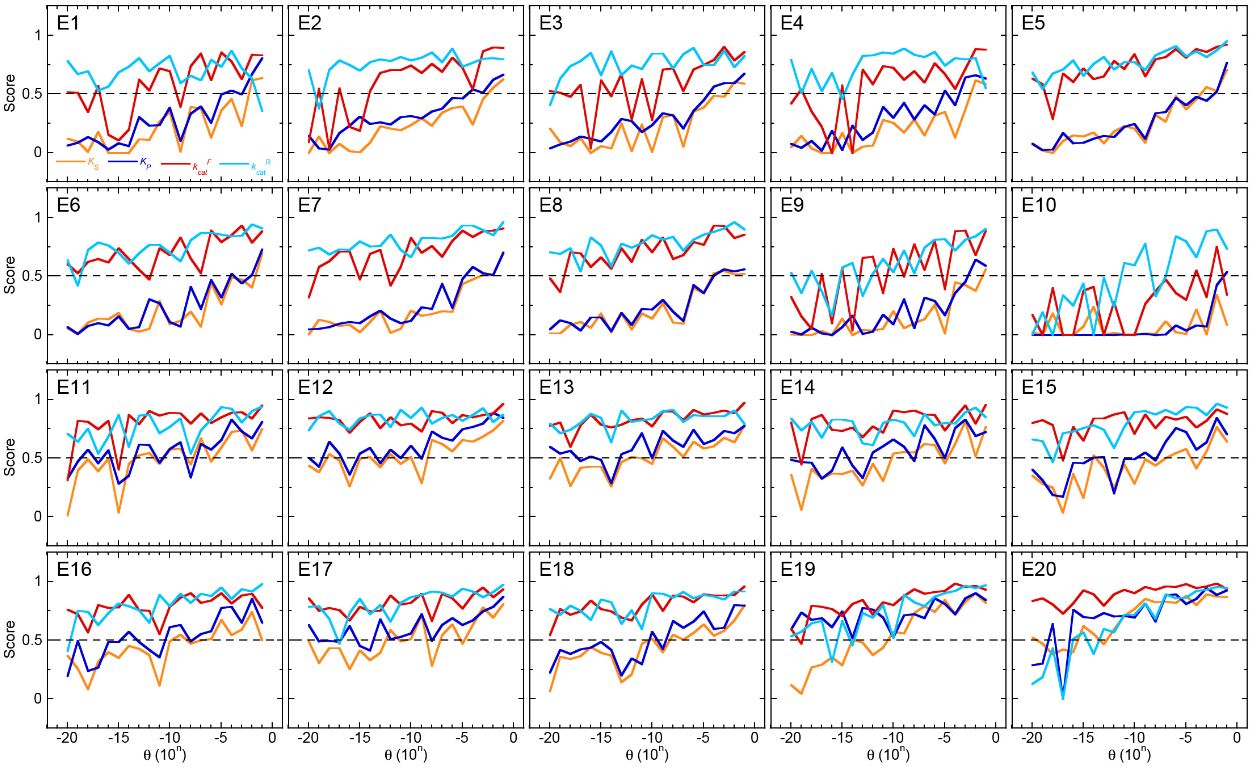


FIGURE S10: **Results for E1-E20 as a function of long simulation time domains.** E1–E20 simulations were performed by varying the time domain ($\theta$). E1-E20 fitting scores for each constant ($K_S$, orange; $K_P$, blue; $k_{cat}^F$, red; $k_{cat}^R$, cyan) were plotted as a function of the $\theta$ ratio (plotted as $10^n$) with error held constant at 3%.

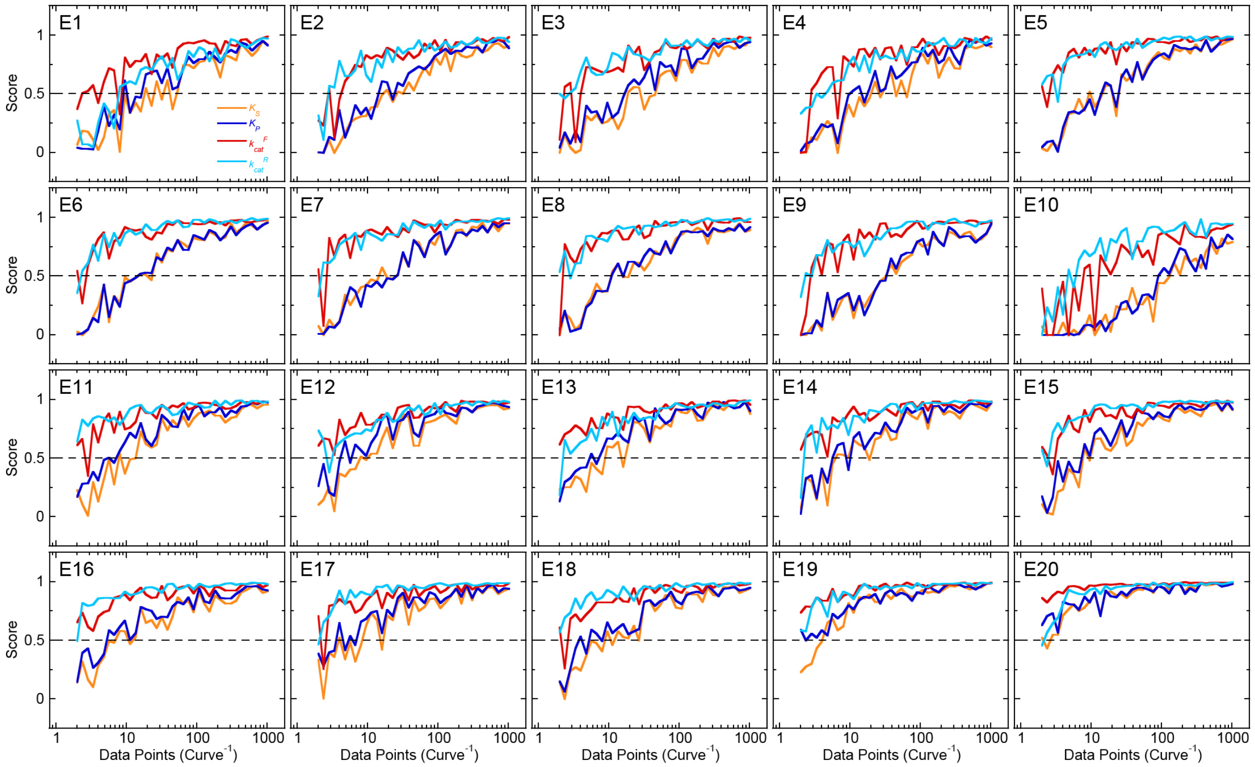


FIGURE S11: **Results for E1-E20 as a function of data point density.** E1-E20 fitting scores for each constant ($K_S$, orange; $K_P$, blue; $k_{cat}^F$, red; $k_{cat}^R$, cyan) were plotted as a function of the number of data points per curve (2-1024 points). Error was held constant at 3% for all simulations.

3. Lambert JH. Observations variae in Mathesin Puram. Acta Helvitica, physico-mathematico-anatomico-botanico-medica. 1758;3:128-68.

4. Euler L. De serie Lambertina Plurimisque eius insignibus proprietatibus. Acta Acad Scient Petropol. 1783;2:29-51.
